# Loss of ovarian hormones modulates the nucleic acid content of circulating extracellular vesicles and skeletal muscle metabolism in response to acute exercise

**DOI:** 10.1101/2025.09.17.676471

**Authors:** V. Puumalainen, A. Maja, T-M. Korhonen, TA. Nissinen, E. Hulkko, J.A. Ihalainen, M. Lehti, S. Karvinen

**Affiliations:** Faculty of Sport and Health Sciences, University of Jyväskylä, Finland; Gerontology Research Center, University of Jyväskylä, Finland; Department of Biological and Environmental Science, University of Jyväskylä, Jyväskylä, Finland; Nanoscience Center, Department of Biological and Environmental Science, University of Jyväskylä, Jyväskylä, Finland; Nanoscience Center, Department of Chemistry, University of Jyväskylä, Jyväskylä, Finland

## Abstract

Loss of ovarian hormones (i.e., menopause) leads to negative effects on metabolic health. While exercise offers significant benefits, it seems to be insufficient to completely reverse these changes. The mechanisms by which exercise conveys the effects throughout the body are still poorly understood. Extracellular vesicles (EVs) are released into circulation during exercise. EVs carry small non-coding RNAs (sRNAs), such as microRNAs (miRs), which are proposed as mediators of the effects of exercise. We have previously shown that the miR response to acute exercise of EVs and HDL is diminished in postmenopausal women with low estrogen levels, which we were able to replicate here also in a rat model. In this study, we examined the effect of loss of ovarian hormones and acute exercise in the sRNA cargo of EV and HDL particles in female rats. We show for the first time that loss of ovarian hormones affects specifically the nucleic acid cargo of circulating EVs. We further show that the estrogen responsive miRs regulate anaerobic glycolytic pathway in skeletal muscle. Finally, we demonstrate that the loss of ovarian hormones leads to higher anaerobic energy production during an acute bout of exercise, implicating an inferior ability to sustain aerobic energy production during exercise.

## 1 INTRODUCTION

Loss of ovarian hormones (i.e. menopause) is associated with several negative health effects, such as increased adipose tissue mass and higher risk for metabolic diseases including type 2 diabetes (Clegg et al., 2017; Juppi et al., 2022). While exercise appears to ameliorate these changes, it cannot completely reverse them (Hyvärinen et al., 2022; Karvinen et al., 2019). We have previously shown that menopause diminishes the acute exercise response of circulating extracellular vesicles (EVs) in postmenopausal women (Karvinen et al., 2023). Yet it is not known how this change in systemic signaling may affect the response to exercise at tissue level.

Withdrawal of ovarian hormones by ovariectomy (OVX) in rodents is widely used to mimic the effects of menopause in the female body. Loss of ovarian hormones in women and female rats is associated with increased body weight and adipose tissue mass, as well as a decline in metabolic health (Asarian & Geary, 2002; Juppi et al., 2022; Lee et al., 2024)). Furthermore, similarly to menopause, the loss of ovarian hormones has been associated with decreased physical activity levels in rodents (Chen et al., 2014; Mäkinen et al., 2023; Pettee Gabriel et al., 2015). We have previously shown that loss of ovarian hormones decreases also the maximal running capacity in female rats (Lee et al., 2024). Accordingly, utilizing OVX model allows for the investigation of the interplay of ovarian hormones and exercise at the tissue level.

While the primary operators of exercise are the skeletal muscles and the cardiorespiratory system, its effects are systemic, influencing the entire body. During recent years, it has been demonstrated that exercise induces release of EVs (Frühbeis et al., 2015; Vechetti et al., 2021; Whitham et al., 2018) and high-density lipoprotein (HDL) particles (Karvinen et al., 2023; Palazón-Bru et al., 2021) into the circulation. EVs and HDL act as information carriers in systemic signaling, carrying for example lipids, proteins and small non-coding RNAs (sRNAs), such as microRNAs (miRs) (Théry et al., 2002; Vickers & Remaley, 2014). However, the role of EVs and HDL in conveying the positive effects of exercise is still poorly understood.

EVs are membrane-bound particles, which are released by all types of cells. EVs have been shown to facilitate cell-to-cell communication (Kalra et al., 2016; Mathivanan et al., 2010) and to act as inter-organ communication mediators, also in response to exercise (Vechetti et al., 2021; Whitham & Febbraio, 2016). Like EVs, also HDL transports sRNAs to recipient cells (Karvinen et al., 2023; Vickers et al., 2011), and this can possibly explain, at least partially, the mechanism by which HDL is able to act as a modulator of recipient cell functions. The sRNA profile of EV cargo differs from that of HDL cargo (Karvinen et al., 2023), suggesting that these communication mediators have complementary but independent mechanisms for sRNA transport. Understanding systemic signaling allows us to uncover the molecular mechanisms underlying the health effects of exercise.

sRNAs are not translated into proteins yet regulate numerous central cellular processes. sRNAs have been shown to respond to exercise (X. Chen et al., 2012; Iguchi et al., 2010; Nielsen et al., 2010; Quinn & Chang, 2016). Of sRNAs, the functions of miRs have been studied most extensively (Gomes et al., 2018). Circulating sRNAs, such as circulating miRs (c-miR), are shown to deliver signals from donor cells to recipient cells both *in vitro* and *in vivo* (Iguchi et al., 2010). Main function of miRs is to negatively regulate protein translation by affecting the stability of messenger RNA (Huntzinger & Izaurralde, 2011). miRs are implicated to play a role for example in metabolism of lipids and glucose (Poy et al., 2007; Rottiers & Näär, 2012), inflammation (Das & Rao, 2022), and coordinating aging-related biological pathways (Inukai & Slack, 2013). Extensive studies have revealed that c-miR profile changes in response to exercise (Barber et al., 2019; F. Li et al., 2020; Nielsen et al., 2014).

Here, we show for the first time that loss of ovarian hormones affects specifically the nucleic acid content of circulating EVs. We also confirm our previous findings, that loss of ovarian hormones diminishes the acute response to exercise in EVs. We further demonstrate that the ovarian hormone responsive miRs coordinate metabolic signaling routes linked to type 2 diabetes. Finally, we show that the anaerobic energy production in skeletal muscle is higher in rats with depleted ovarian hormones, suggesting an inferior capacity to maintain aerobic energy production in response to acute exercise. Our results highlight the role of ovarian hormones in the acute exercise response and possibly in mediating the health benefits of exercise.

## 2 MATERIALS AND METHODS

### 2.1. Animals

The study sample consisted of 40 female Wistar Hans rats, which correspond to a representative subset of study design reported previously by us (Figure 1 a,b) (Lee et al., 2024). Rats were purchased from Envigo (Indianapolis, IN, United States, bred and shipped from the Netherlands), and arrived at the animal facilities of University of Jyväskylä at the age of 13–15 weeks. All rats were housed in pairs in an environment-controlled facility (12/12 h light–dark cycle, 22°C) and received water and estrogen-free rodent feed (2019X, Envigo, Indianapolis, United States) *ad libitum* upon the arrival. Half of the rats (n=20) underwent ovariectomy (OVX), resulting in the loss of ovarian hormones and the other half (n=20) was predisposed to sham surgery without the removal of ovaries. The groups were matched for body mass and maximal running capacity. To ensure that the rats were fully grown adults prior to the surgeries, the surgeries were performed at the age of 7 months (Figure 1a) (Yousefzadeh et al., 2020). The OVX and sham groups were further divided into two groups, one performing maximal running test (max) to induce an acute exercise stimulus, and the other serving as a control. Hence, the resulting groups were sham/control, sham/max, OVX/control and OVX/max (Figure 1b). Rats in the sham group were euthanized at proestrus (determined by cytology sample) to ensure that they were in the highest possible systemic estrogen state at the time samples were collected (Lee et al., 2024).

**FIGURE 1.**
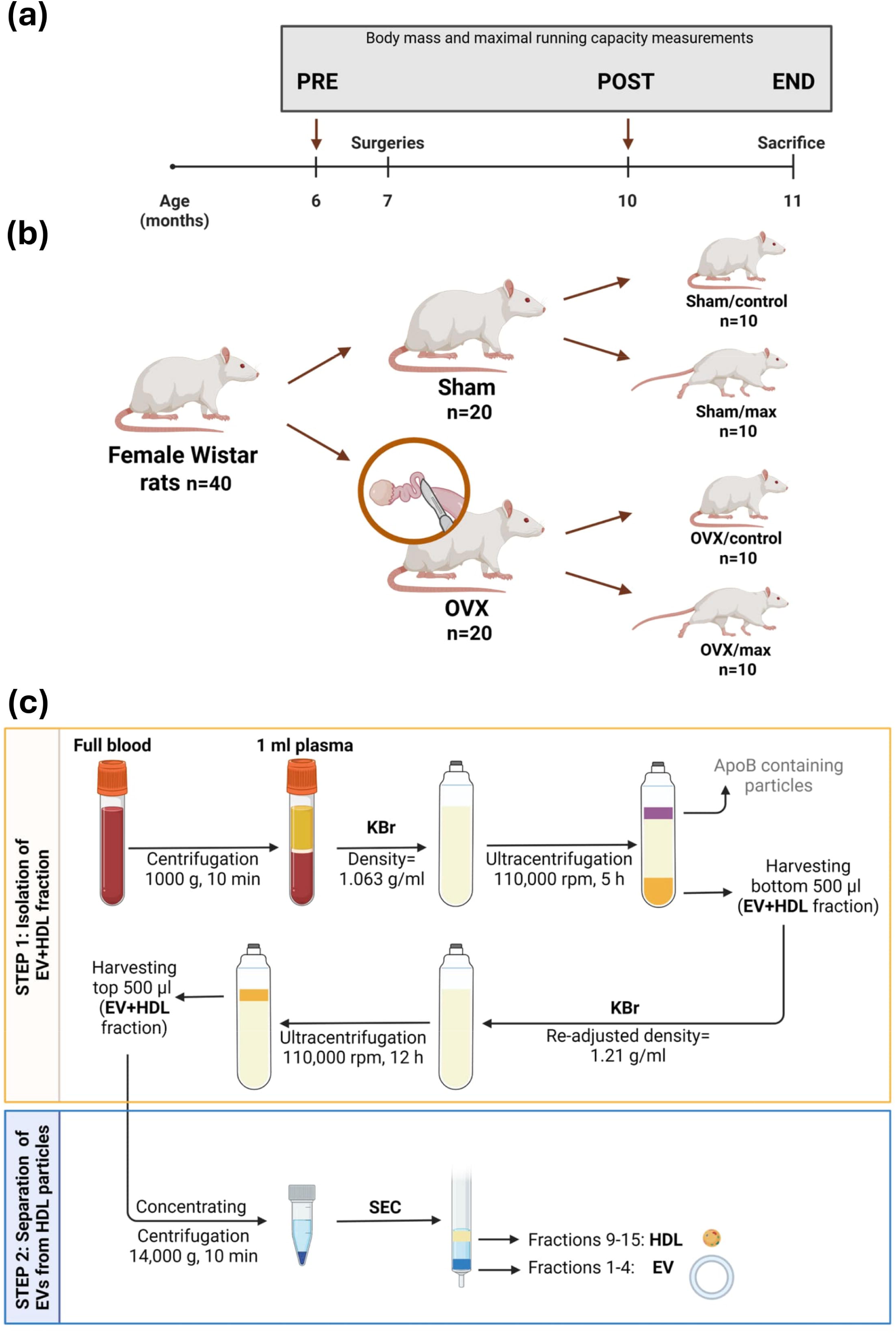
Schematic representation of the study design. (a) Timeline of the study. Rats were matched for body weight and maximal running capacity before dividing them into groups. (b) 40 female Wistar rats were divided to two groups: ovariectomy (OVX), and sham surgery without the removal of ovaries (sham). Groups were further divided so that half of the rats performed a maximal running test to induce an acute exercise stimulus. (c) Flowchart of EV and HDL particle isolation protocol. STEP 1: EVs and HDL were isolated based on density, STEP 2: EVs were separated from HDL particles based on size. OVX=ovariectomy, EV=extracellular vesicle, HDL=high-density lipoprotein, KBr=potassium bromide, ApoB=apolipoprotein B, SEC=size-exclusion chromatography.

### 2.2. Ethics statement

The study has received approval from national Project Authorization Board (ELLA, Finland, permit number ESAVI/4209/2021) and was conducted in accordance with the “Principles of Laboratory Animal Care” (NIH publication #85–23, revised in 1985) and the European Commission Directive 2010/63/EU.

### 2.3. Maximal running capacity test

Rats were tested for their maximal running capacity similarly as described in (E. Lee et al., 2024). Briefly, a speed-ramped treadmill running test (15° slope, initial velocity of 10 m/min, increased 1 m/min every 2 min) was performed PRE and POST the intervention, and for max groups immediately before euthanasia (Figure 1a). First, rats were habituated to running on the treadmill with three different sessions lasting for 10 min with a low velocity (<10 m/min). Maximal running test was repeated three times with at least 1 day of recovery in between. The best result of the three trials [maximal running distance (m)] was considered the maximal running capacity, except for the test performed immediately before euthanasia for the max groups, where the test was performed only once.

### 2.4. Blood sampling

At the end of the study, rats were fasted for 2h and euthanized with carbon dioxide followed by heart puncture. Tissue samples were collected, weighed and snap frozen in liquid nitrogen and stored at -80°C until further analyses. Of the adipose tissue deposits, only retroperitoneal adipose tissue was collected and weighed because of its more distinct location and uniform composition compared to visceral (i.e., omental) and/or ovarian adipose tissue deposits. Plasma (EDTA) was separated from the whole blood by 15-minute incubation at RT, followed by centrifugation (1,000 × g, 10 min at RT). Plasma was stored as 200 μl aliquots at -80°C.

### 2.5. Isolation of EVs and HDL

The isolation of EV and HDL was carried out similarly as described before (Karvinen et al., 2023). The workflow of isolation and separation of HDL particles and EVs from plasma is depicted in Figure 1c.

EV and HDL -containing fractions were isolated by sequential ultracentrifugation, in which the density was adjusted by potassium bromide (KBr) to separate lipoproteins by density difference (Havel et al., 1955). Plasma samples (1,000 µl) were thawed, and their volume was adjusted to 2 ml with sterile filtered (0.2 µm) PBS and density to 1.063 g/ml and ultracentrifuged at 110,000 rpm at +4°C, (Beckman, 110 TLA rotor) for 5 h in 3.2 ml ultracentrifuge tubes (Beckman Coulter, 362333). Bottom 500 µl was collected, volume adjusted to 2 ml and density readjusted to 1.21 g/ml and ultracentrifuged (110,000 rpm, +4°C) for 12 h. The EV and HDL containing top 500 µl fraction was collected and stored at - 80°C. Ultracentrifugation results in fraction containing both EVs and HDL, and therefore size-exclusion chromatography (SEC) was performed to separate EVs from HDL. Prior to SEC, ultracentrifuged HDL and EV fraction was concentrated by centrifugation (14,000 × g, +4°C, Heraeus Fresco 17 centrifuge) with Amicon® Ultra centrifugal filters (100 kDa, cat. UFC5100, Merck) according to the manufacturer’s instructions. Concentrates were stored at - 80°C until SEC.

EVs and HDL were separated by SEC (IZON qEV1, 35nm, IC1-35). Concentrate containing HDL and EVs was thawed on ice. SEC column was operated as instructed by the manufacturer at room temperature and at upright position. To reach recommended sample volume of 1 ml, sterile, filtered (0.2 µm) PBS (pH 7.4) was added to the concentrated sample. 1 ml of concentrated sample was loaded on to the column and 4.7 ml void volume was discarded. Then, 16 fractions were collected, 700 µl each. EVs were collected to fractions 1-4 according to the manufacturer’s instructions. From SEC fractions, EVs and HDL were concentrated the same way as prior to SEC.

Throughout the protocol, the EV and HDL containing samples were kept on low-binding tubes (Low protein binding microcentrifuge tubes, Thermo Scientific, Cat no. 90410) when applicable to reduce the loss of particles by adhesion to the microcentrifuge tubes.

### 2.6. Validation of EV and HDL isolation

EV zone was located in fractions 1-4 and HDL zone was located between SEC fractions 9-15, which was confirmed by bicinchoninic acid assay (BCA) (Pierce Biotechnology, Rockford, IL) with automated instrument (KoneLab, Thermo Scientific, Vantaa, Finland), dot blot, and electron microscopy (EM) (Figure S1).

The amount of protein in the EV samples could not be visualized by traditional WB (*data not shown*), so a dot blot assay was performed by pipetting 1 µl of sample directly onto nitrocellulose membrane. For total protein visualization, Ponceau S staining was used, whereas for EVs and HDL, antibodies for CD63 (1:500; cat. #ab59479, Abcam), APOA1 (1:10,000; cat. #ab52945, Abcam), and TSG101 (undiluted; cat. #ab125011, Abcam) were used. CD63 is a transmembrane protein, enriched in EVs (Escola et al., 1998), TSG101 a cytosolic protein expressed in EVs, whilst APOA1 is enriched especially in HDL, but is detected also in EVs (Karimi et al., 2018). Total protein and dot blot confirmed that HDL was enriched in fractions 9-15.

Samples containing EVs and HDL separately (the fractions after SEC) were prepared for EM as described previously (Karvinen et al., 2020; Puhka et al., 2017). Briefly, EV and HDL samples were loaded on 200 mesh grids, fixed with 2% paraformaldehyde solution (PFA), stained with 2% neutral uranyl acetate, and embedded in a uranyl acetate and methyl cellulose mixture (1.8/0.4%). Samples were viewed with transmission EM using a Jeol JEM-1400 (Jeol Ltd., Tokyo, Japan) operating at 80 kV. Images were taken with a Gatan Orius SC 1000B CCD-camera (Gatan Inc., United States).

### 2.7. Raman spectroscopy

The molecular composition of the EV cargo was studied via Raman spectroscopy from pooled, representative samples of the study groups. For this purpose, EVs were isolated from plasma using SEC similarly as described in 2.2. Isolation of EVs and HDL. EV samples were analyzed using Raman spectroscopy (Nicolet DXR Raman Microscope) equipped with a diode-pumped solid-state laser operating at 532 nm following a previously described protocol (Gualerzi et al., 2019). Briefly, 5 µl of EV suspension was deposited on a gold slide. All the measurements were performed on the air-dried drop with 50x objective, 900 lines/mm diffraction grating, 700 µm spot size, and confocal mode (50 µm pinhole) in the spectral range of 400−3581 cm^−1^. For each group, five measurements were collected and used for the analysis.

The Raman spectra were processed using Matlab by fitting with two linear lines to subtract the background from the signal. Thereafter, the signal was normalized to the highest lipid peak. To calculate the nucleic acid to lipid (NA/L) and protein to lipid (P/L) ratios as described previously (Mihály et al., 2017), we first calculated the average of each molecular class (nucleic acid band 720-800 cm^-1^, amide I protein band (1600-1690 cm^-1^) and lipid-related band (2750-3040 cm^-1^)). Unfortunately, we were unable to assess the molecular cargo of HDL by Raman spectroscopy.

### 2.8. Small RNA extraction and sequencing

Before RNA extraction, the SEC fractions containing EVs and HDL particles were concentrated similarly as described above. RNA was extracted using miRNeasy Serum/Plasma Kit (Qiagen, Germany, Cat no. 217184) according to the manufacturer’s instructions. Thereafter, 10 µl of RNA isolate was shipped to Novogene (Germany) for exosomal small RNA library preparation and sequencing.

### 2.9. Raw sRNA data processing and alignment

Clean reads were received from Novogene in FASTA format. Reads shorter than 20 nucleotides were excluded. For the analysis of miRs, the reads were trimmed to 22 bp using a FastX-Toolkit ((Gordon & Hannon, 2010, available online at: http://hannonlab.cshl.edu/fastx_toolkit). The alignment was done using Bowtie (Langmead et al., 2009). No reverse complement -option was used. Only one best alignment for a read was used as output, even if there were multiple possible alignments. miRs were aligned to miRBase version 22 (Kozomara et al., 2019), ribosomal RNA (rRNA) -derived sRNAs (rDRs) were aligned to the full set of rat rRNA (downloaded from RNAcentral in February 2025), and transfer RNA (tRNA) -derived sRNAs (tDRs) were aligned to the high confidence tRNA gene set from GtRNAdb (Chan & Lowe, 2016). For miR sequences, 2 mismatches were allowed. For tDR sequences, 1 mismatch was allowed, and for rDR sequences, no mismatches were allowed.

### 2.10. TIGER pipeline analysis of sRNA

The sRNA species in EV and HDL particles were analyzed with “Tools for Integrative Genome analysis of Extracellular sRNAs (TIGER) pipeline (Allen et al. 2018). This analysis categorizes the sRNA species by their origin by utilizing genome and database alignments (Figures3-4).

### 2.11. Differential expression analysis on sRNAs

Differential expression (DE) analyses were done as described earlier (Karvinen et al. 2023) using DESeq2 R-package in R-program (v 4.3.1) for EV and HDL samples (*n*=10 per group each). Samples with fewer than ten sRNA (miR, rDR, or tDR) counts in at least seven samples in a subgroup were excluded from the analysis.

### 2.12. miR target analysis and signaling pathway heatmap

To elucidate the possible functional roles of EV- and HDL-carried miRs that were differentially expressed in response to exercise, a miR target analysis was done via miRNet (https://www.mirnet.ca/) (Chang et al., 2020). In EV miR analyses the tissue was set to exosomes, whereas in HDL miR analyses, the tissue was unspecified. Minimal pathway analysis was used throughout. The possible interactions of miRs and mRNAs were examined by miRPath v3.0 (Vlachos et al., 2015). In miRPath analysis, microT threshold was set to 0.08, and p-value of 0.06 was used to avoid excluding several essential exercise-related miR signaling pathways. Central pathways to energy production were analyzed further, and target proteins of differentially expressed miRs were examined from these pathways.

### 2.13. Western blot (WB) analysis of the miR target proteins

Western blot (WB) analysis was done to measure the target protein levels of the differentially expressed miRs detected from the EV and HDL fractions. The analyzed proteins were mammalian target of rapamycin (mTOR), insulin receptor substrate 2 (IRS2) and pyruvate kinase L/R (PKLR), which were chosen based on the miR target analysis via miRNet (section 2.9 miR target analysis and signaling pathway heatmap). It has been noted that EVs released after exercise tend to localize to liver tissue (Whitham et al., 2018), and therefore the expression of target proteins in liver was investigated. Given its role in exercise, it was reasoned to analyze also muscle protein levels. In addition, both tissues are crucial for energy metabolism. (Whitham et al., 2018).

First, the liver and gastrocnemius muscle samples (30 mg) were homogenized in buffer solution [20 mM HEPES (pH 7.4), 1 mM EDTA, 5 mM EGTA, 10 mM MgCL_2_, 100 mM β-glycerophosphate, 1 mM Na_3_VO_4_, 2 mM DTT, 1% NP-40, 0.2% C_24_H_39_O_4_Na, and 3% protease and phosphatase inhibitor cocktail (Thermo Scientific, Cat no. 78443)] in TissueLyser II (Qiagen) for 2 × 2 min, 30 Hz. Next, the homogenates were kept on a sample rocker for 30 minutes at +4°C, and centrifuged at 10,000 × g at +4°C for 10 minutes. Total protein was determined by bicinchoninic acid (BCA) assay with an automated instrument (KoneLab, Thermo Scientific, Vantaa, Finland) and diluted to 2 µg/µl. Laemmli buffer was added to the samples and the samples were heated for 10 minutes to linearize the proteins. Separation was done by SDS-PAGE for 30-40 min at 270 V on Criterion electrophoresis cell (Bio-Rad Laboratories, Richmond, CA) on 4-20% gradient gels (Criterion™ TGX Stain-Free™ Precast Gel, Bio-Rad, Cat no. 5678094). Proteins were transferred onto nitrocellulose membrane in Turbo blotter (Trans-Blot Turbo Blotting System, 170-4155, Bio-Rad Laboratories). Thereafter, the membrane was blocked for 2 h at room temperature in a commercially available blocking buffer (Intercept® (PBS) Blocking Buffer, LI-COR Biosciences, Cat no. 927-70001). Then the membrane was incubated overnight at 4°C in primary antibody solution to measure target proteins: mTOR (2983S, Cell Signaling), IRS2 (ab134101, Abcam) and PKLR (ab137787, Abcam), all at dilution 1:1,000 in 1:1 blocking buffer and TBS. After the incubation, membrane was washed in TBS-T and incubated in suitable secondary antibody diluted in 1:1 blocking buffer and TBS-T for 2h at room temperature and washed again in TBS-T. Fluorescence images were obtained with ChemiDoc MP™ Imaging System together with Image Lab™ Touch software (version 2.4.0.03, Bio-Rad Laboratories). Proteins were quantified using Image Lab software (version 6.1.0). The blot lanes were first normalized to blot average, and the individual bands were subsequently normalized to corresponding lane averages. The results are expressed as arbitrary units (AU).

### 2.14. Lactate dehydrogenase enzyme activity measurement

To assess the anaerobic energy production in skeletal muscle, the same gastrocnemius muscle homogenates as utilized for the WB analysis were subjected to lactate dehydrogenase (LDH) analysis (Thermo Fisher, Cat no. 981906). The muscle homogenates were diluted 1:20 into milliQ H_2_O and the LDH activity was measured from single samples (n=10/group) via Indiko Plus (Thermo Fisher Scientific, Vantaa, Finland).

### 2.15. Statistics

Statistical analyses were conducted using IBM SPSS (version 28.0.1.1). The normality was assessed by Shapiro-Wilk test followed by Levene’s test. When data was normally distributed, group comparisons were made with Student’s *t*-test, and when the data did not meet the criteria for normal distribution, Mann-Whitney U-test was used. Two-way ANOVA was used for assessing the main effects of OVX, acute exercise and their interaction. A *p*-value <0.05 was regarded as statistically significant.

## 3 RESULTS

### 3.1. The loss of ovarian hormones increased body fat mass and reduced maximal running capacity

As OVX surgery is known to increase body mass and decrease physical activity level, the rats were matched for their body mass and maximal running capacity before dividing them into sham and OVX groups (PRE, p≥0.196, Table 1, Figure 1). Furthermore, the sham and OVX groups were divided into control and max groups, matched for body mass and maximal running capacity (PRE, p≥0.459, Table 1, Figure 1.).

**TABLE 1.** Characteristics of the study groups.

|  | Sham/control<br>n=10 | Sham/max<br>n=10 | OVX/control<br>n=10 | OVX/max<br>n=10 | p-value<br>(sham/OVX) | p-value<br>(sham/control<br>vs. sham/max) | p-value<br>(OVX/control<br>vs. OVX/max) |
| --- | --- | --- | --- | --- | --- | --- | --- |
| Body mass (g) |  |  |  |  |  |  |  |
| PRE | 276.3 (34.0) | 280.3 (37.2) | 298.4 (42.2) | 286.8 (23.8) | 0.196 | 0.805 | 0.459 |
| POST | 312.4 (44.2) | 309.1 (41.2) | 364.2 (50.3) | 359.7 (35.5) | **<0.001** | 0.865 | 0.820 |
| END | 315.5 (46.6) | 315.0 (48.4) | 373.0 (59.4) | 363.0 (36.7) | **0.001** | 0.981 | 0.657 |
| Total muscle mass† (g) | 2.58 (0.29) | 2.67 (0.30) | 2.75 (0.27) | 2.67 (0.23) | 0.311 | 0.473 | 0.472 |
| Total muscle mass† (% of body mass) | 0.82 (0.07) | 0.85 (0.06) | 0.75 (0.08) | 0.74 (0.03) | **<0.001** | 0.303 | 0.656 |
| Retroperitoneal fat mass (g) | 5.56 (2.57) | 5.51 (2.94) | 9.82 (4.33) | 10.71 (2.46) | **<0.001** | 0.971 | 0.580 |
| Uterine mass (g) | 0.69 (0.17) | 0.71 (0.10) | 0.13 (0.02) | 0.15 (0.08) | **<0.001<sup>b</sup>** | 0.729 | 0.529<sup>b</sup> |
| Liver mass (g) | 9.15 (0.97) | 9.16 (1.30) | 8.70 (1.83) | 7.96 (0.70) | **0.047** | 0.982 | 0.243 |
| Maximal running capacity (m) |  |  |  |  |  |  |  |
| PRE | 395.3 (116.8) | 378.6 (178.6) | 363.1 (159.7) | 373.3 (153.0) | 0.694 | 0.529<sup>b</sup> | 0.886 |
| POST | 410.8 (114.7) | 395.2 (164.5) | 327.7 (176.4) | 301.0 (87.9) | **0.009<sup>b</sup>** | 0.631<sup>b</sup> | 0.853<sup>b</sup> |
| END |  | 275.3 (86.5) |  | 238.8 (77.7) | 0.334 |  |  |

\*b=Mann-Whitney U-test, bold= $p < 0.05$ . †= Calculated sum of average muscle mass (left and right) hindlimb muscles (including gastrocnemius, extensor digitorum longus, tibialis anterior, and soleus) was used to represent the total muscle mass.

Within-group (OVX or sham) comparisons confirmed that the subgroups (max vs. control) did not differ in the studied variables (p≥0.243, Table 1). However, as expected based on the previous study with larger study cohort (Lee et al., 2024) of which the study design reported here is a subset, after the intervention sham vs OVX comparisons revealed significant differences in several variables (Table 1). The OVX group had higher body mass and retroperitoneal fat mass, and lower relative total muscle mass (% of body weight) than the sham group (all p<0.001), although absolute total muscle mass (g) was similar between the groups (p=0.311, Table 1). The uterine mass was significantly lower in OVX than in sham, confirming successful ovariectomy surgeries (p<0.001, Table 1).

### 3.2. The loss of ovarian hormones changed the nucleic acid-to-lipid ratio in EVs in response to acute exercise

The enrichment of EVs and HDL into separate fractions was confirmed by BCA, dot blot, and EM (Figure S1). We examined the EV molecular cargo of pooled, representative samples of each study group via Raman spectroscopy. Furthermore, we calculated the nucleic acid to lipid (NA/L) and protein to lipid (P/L) ratios.

The Raman spectra exhibited consistent peak patterns across all groups, suggesting uniform quality among the samples (Figure 2a). However, despite the Raman spectra similarity between groups, the spectral subtraction graphs showed differences between the groups (Figure 2b). We observed a significant effect of acute exercise and an interaction of OVX and acute exercise in nucleic acid to lipid (NA/L) ratio (p≤0.039, Figures 2c-d). OVX/max had a higher NA/L ratio compared with OVX/control group (p=0.001, Figures 2c-d). There were no significant findings from protein to lipid (P/L) ratio (p≥0.166, Figures 2c-d). Our results suggest that specifically the nucleic acid cargo of EVs is affected by the loss of ovarian hormones and acute exercise.

**FIGURE 2.**
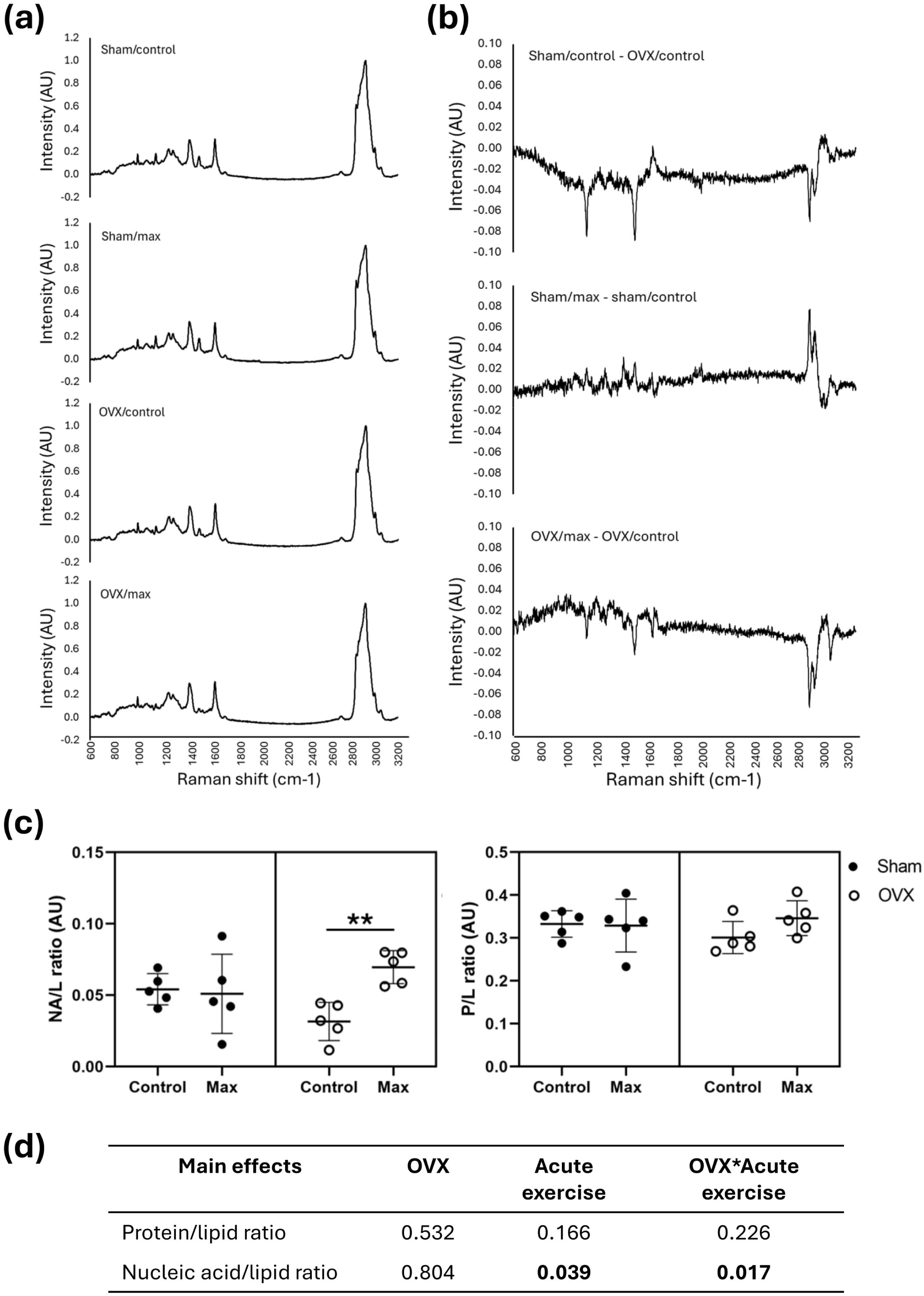
Loss of ovarian hormones (induced by OVX) changes specifically the nucleic acid cargo of EVs in response to exercise. (a) Full Raman spectra exhibit consistent peak patterns across the groups. (b) Subtraction spectra display differences between groups. (c) OVX and acute exercise affect significantly the nucleic acid-to-lipid (NA/L) ratio, whereas protein to lipid (P/L) ratio did not display changes. (d) Main effects of OVX, acute exercise and the interaction of OVX and acute exercise on P/L and NA/L ratio (p-values). Acute exercise and the interaction of OVX and acute exercise (OVX*Acute exercise) affect significantly the NA/L ratio, whereas no effects were seen on the P/L ratio. OVX=ovariectomy, NA=nucleic acid, L=lipid, P=protein.

### 3.3. miRs displayed a greater portion of host sRNA cargo in EV than HDL particles

Since the nucleic acid cargo of EVs was the most affected by OVX and acute exercise, we continued by examining the sRNA species of EV and HDL particles. There was an average of 10.6 (8.4 – 15.8) million raw reads/sample in EV samples, whereas HDL samples had an average of 10.0 (6.4 – 14.3) million raw reads/sample. After adapter trimming, the average sRNA read count was 8.2 million in EVs, and 8.4 million in HDL, of which miRs composed 144,000 in EVs and 103,000 in HDL.

We began our analysis by inspecting the sRNA origin in EV and HDL particles. In both particles, major portion of the sRNAs were either unmapped or found to be too short for mapping (<16 nucleotides) (Figure 3a). Of the mapped sequences, a greater share was mapped to nonhost origin (26 – 36%), than to host origin (10 – 13%) (Figure 3a). The relative sRNA abundances were similar between groups, and EV and HDL particles also shared similarities (Figure 3b). Majority of EV nonhost sRNA sequences were mapped to rDR (46 – 48%), and to more than one category (37 – 40%) (Figure 3b). In HDL, most of nonhost sRNA sequences were mapped to same categories as in EV particles, but more were mapped to more than one category than to rDR (43 – 48%, and 36 – 42%, respectively, Figure 3b). Of the sRNA host sequences, majority were mapped to rDR both in EV and HDL particles (50 – 55%, and 49 – 57%, respectively, Figure 3c). In EVs, the second most common category was miRs, constituting 19 – 30% of all sRNA species. The abundance of miRs in EVs was lowest in sham/control group (19%), whereas other groups did not portray differences in miR abundance (29% in sham/max and OVX/control, and 30% in OVX/max, Figure 3c). In HDL, miR abundance in host sequences was higher in the sham groups (27 and 28% in sham/control and sham/max, respectively), than in the OVX groups (23% in both OVX/control and OVX/max) (Figure 3c).

**FIGURE 3.**
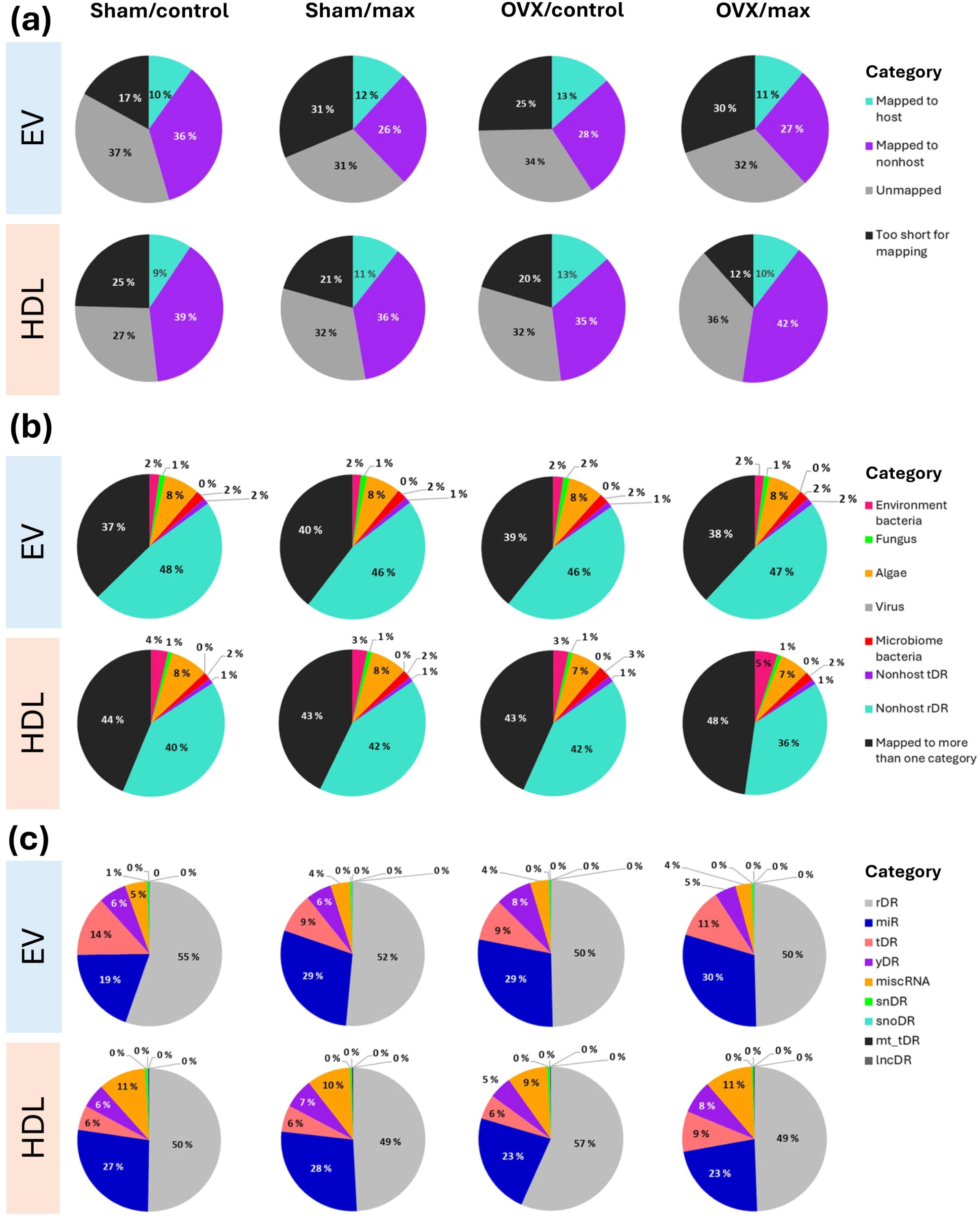
EV and HDL particle sRNA content in the study groups. (a) Majority of the mapped sRNAs of EV and HDL particles originated from nonhost. (b) Of the mapped sRNAs, most in EV and HDL particles were mapped to nonhost ribosomal RNA-derived sRNAs (rDRs). (c) Of the host sRNAs, the majority were mapped to rDR in both EV and HDL particles. rDR= ribosomal RNA (rRNA) -derived sRNA, miR=microRNA, tDR=transfer RNA (tRNA) -derived sRNA, yDR=yRNA, miscRNA=miscellaneous RNA, snDR=small nuclear RNA (snRNA) -derived sRNA, snoDR=small nucleolar RNA (snoRNA) -derived sRNA, mt_tDR=mitochondrial tDR, lncDR=long non-coding (lncRNA) -derived sRNA.

The top ranked sRNA sequencies in EVs and HDL were similar between groups, with the major class being nonhost rDR (Figure 4a). In EVs, miRs constituted a larger proportion of sRNA classes compared to HDL particles (Figure 4a). Most of the top 100 nonhost sequences in EVs were mapped to rDR, while in HDL, the majority were environment-derived sRNAs (Figure 4b). Most of the top 100 host sequences of EV sRNAs were mapped to miR, rDR, and tDR. In HDL, the top 100 sRNA classes were the same as in EVs, with the distinction that tDRs constituted a smaller proportion than in EVs (Figure 4c).

**FIGURE 4.**
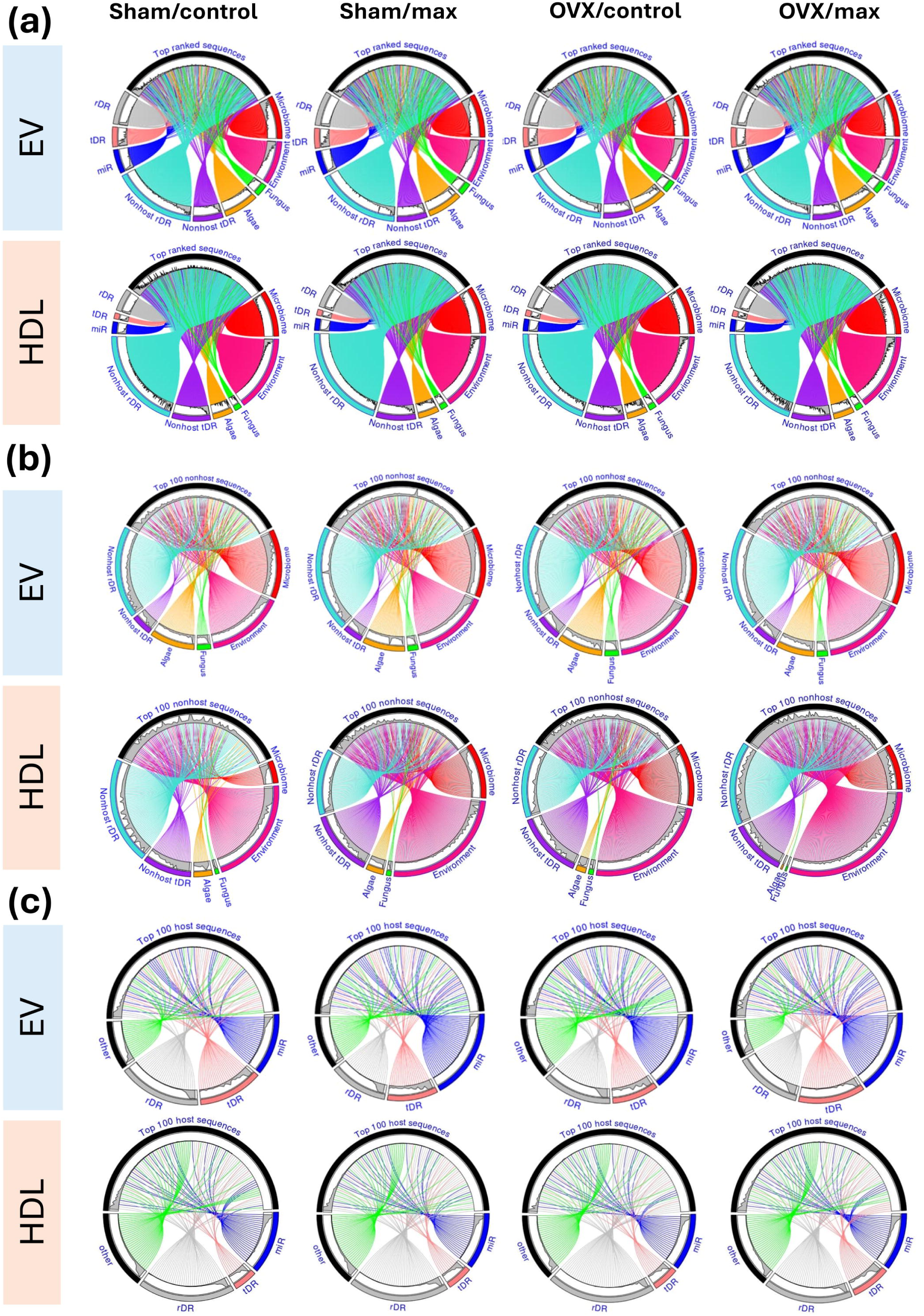
The top sRNA species of EV and HDL particles displayed minor differences. (a) Top ranked species in EV and HDL particles in each study group. (b) Top 100 ranked nonhost sRNA species in EV and HDL particles in each study group. (c) Top 100 host sRNA species in EV and HDL particles in each study group. rDR= ribosomal RNA (rRNA) -derived sRNA, tDR=transfer RNA (tRNA) -derived sRNA, miR=microRNA.

### 3.4. The loss of ovarian hormones blunted the acute exercise-induced miR response of EVs

We examined the 20 most common miRs in EV and HDL particles and observed 17 common miRs in EVs and 16 in HDL particles between the study groups (Figures 5a,e). The counts of all observed miRs are presented in Table S1.

**FIGURE 5.**
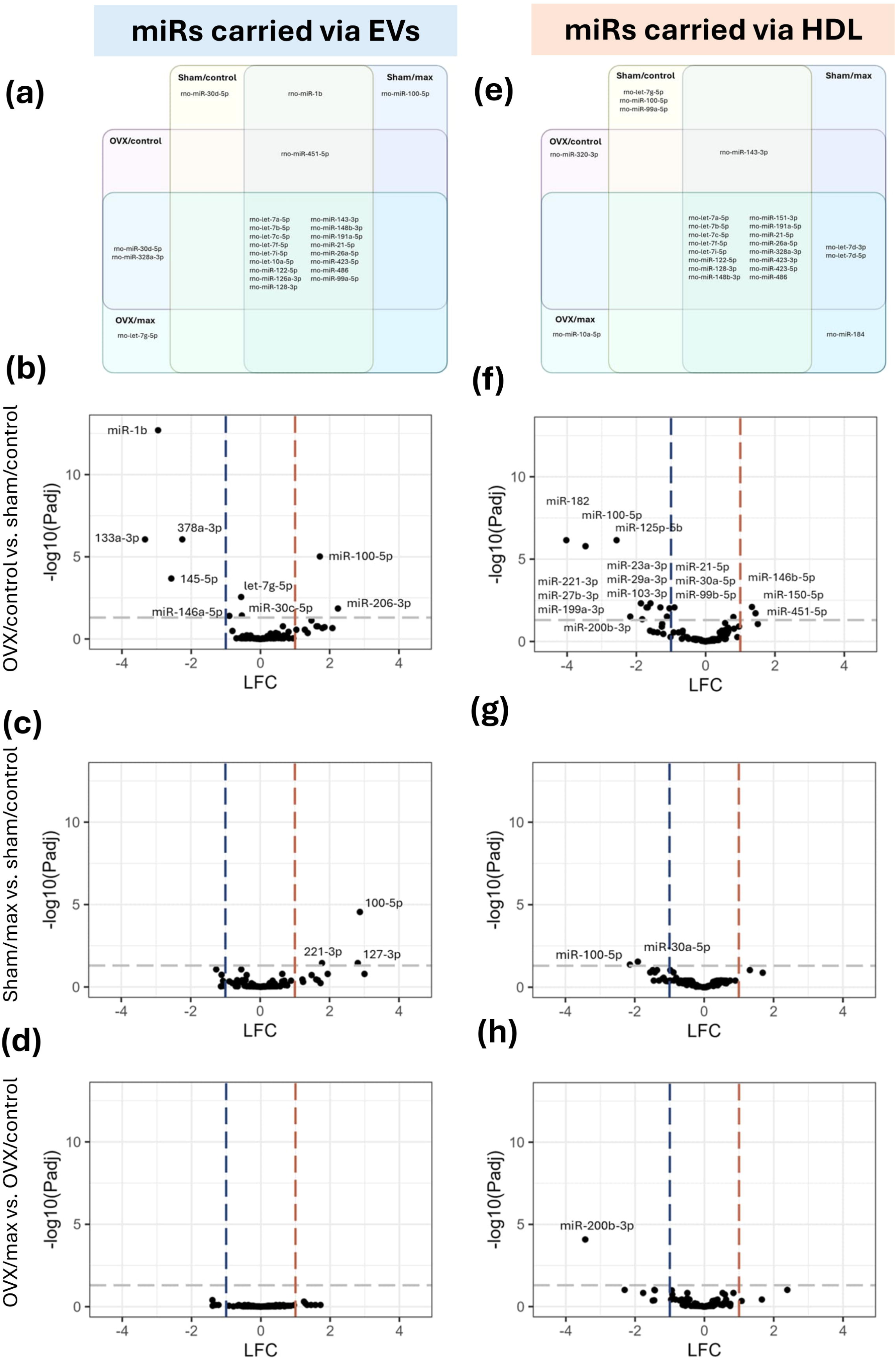
The miR cargo of EV and HDL particles responds to an acute bout of exercise differently. (a) Twenty most common miRs present in EV (reads). (b-d) Volcano plots showing group comparisons of differential expression of miRs in EV particles. (e) Twenty most common miRs present in HDL (reads). (f-h) Volcano plots showing group comparisons of differential expression of miRs in HDL particles. OVX=ovariectomy, miR=microRNA, LFC=log fold-change.

We further examined the miR levels in EVs based on ovarian hormone status (OVX/control vs. sham/control), and the levels of nine miRs were significantly different between the groups (p<0.05, Figures 5b,f, Table S2). Of these, miR-1b, miR-133-2p, miR-378a-3p, miR-30c-5p, let-7g-5p and miR-145-5p were downregulated (p<0.05, Figure 5b, Table S2), whereas miR-100-5p and miR-206-3p were upregulated (p<0.05, Figure 5b, Table S2) in OVX. Furthermore, miR-1b, let-7g-5p, and miR-100-5p were among the twenty most common miRs in EVs, indicating a high biological relevance (Figure 5a, Table S1). When examining the effect of acute exercise in sham groups, there were three significantly upregulated miRs in EVs: miR-100-5p, miR-127-3p and miR-221-3p (p<0.05, Figure 5c, Table S2). Of the significantly different miRs, miR-1b, miR-100-5p and let-7g-5p were also among the twenty most common miRs (Figure 5a, Table S1). Excitingly, no differences between the OVX groups (OVX/max vs. OVX/control) were observed in EVs (p>0.390, Figure 5d).

When examining the effect of ovarian hormones in HDL particles, there were 16 significantly different miRs: miR-146b-5p, miR-150-5p, and miR-451-5p were upregulated (p<0.05, Figure 5f, Table S3) and miR-21-5p, miR-23a-3p, miR-27b-3p, miR-29a-3p, miR-30a-5p, miR-99b-5p, miR-100-5p, miR-103-3p, miR-125b-5p, miR-182, miR-199a-3p, miR-200b-3p, and miR-221-3p downregulated (p<0.05, Figure 5f, Table S3) in OVX. Of the significantly different miRs, miR-21-5p, 99b-5p and 100-5p were also among the twenty most common miRs (Figure 5e, Table S1). When examining the effect of acute exercise in sham groups, there were two significantly downregulated miRs in HDL particles: miR-30a-5p and miR-100-5p (p<0.05, Figure 5g, Table S3). Interestingly, when comparing HDL miRs in OVX groups, miR-200b-3p was significantly downregulated (p<0.001, Figure 5h, Table S3) in the exercised (max) group. However, miR-200b-3p was not among the twenty most common miRs indicating potentially minor physiological relevance (Table S1).

When examining the rDR and tDR species, we observed three significantly lower rDRs in HDL particles in OVX/max when comparing to OVX/control, all of which originated from small subunit ribosomal (SSU) rRNA (Figure S2, Table S5). There were no significant findings in rDRs carried via EVs (Figure S2, Table S4), nor in HDL in other group comparisons (Figure S2, Table S5). In tDRs, there were two significantly higher species in EVs when comparing control groups of OVX and sham; tRNA-Met-CAT-3-1 and tRNA-Met-CAT-1-1. Only tRNA-Met-CAT-3-1 was significantly higher in sham/max vs. sham/control (Figure S3, Table S6). There were no significant differences in tDRs between the OVX groups in EVs or in the tDRs carried via HDL particles between any of the studied groups (Figure S3, Table S7).

### 3.5. The exercise-responsive miR cargo of EVs and HDL may affect energy metabolism in skeletal muscle through target proteins

To study the ovarian hormone-responsive miRs’ functional relevance, a miR-mRNA pathway analysis was carried out with miRPath v3.0 (Vlachos et al., 2015). The analysis was performed to all miRs found to be differently expressed between OVX/control and sham/control, i.e., for nine EV-carried, and 16 HDL-carried miRs (Figures 5b,f, Table S2). Central pathways to energy production were analyzed further, and target proteins of differentially expressed miRs were examined from these pathways.

Analysis of EV miRs, using Kyoto Encyclopedia of Genes and Genomes (KEGG) analysis with pathways union option showed two pathways to be significantly regulated by three of the studied EV-carried miRs, involving type 2 diabetes pathway (Figure 6a, Table S8). Type 2 diabetes signaling pathway regulation by the three miRs was mediated through the following target proteins: mTOR, PKLR, IRS2, and calcium voltage-gated channel subunit alpha1 D (CACNA1D) (Figure 6a). Similar analysis of ovarian hormone-responsive HDL miRs revealed seven pathways to be targeted by four of the studied miRs (Table S9).

**FIGURE 6.**
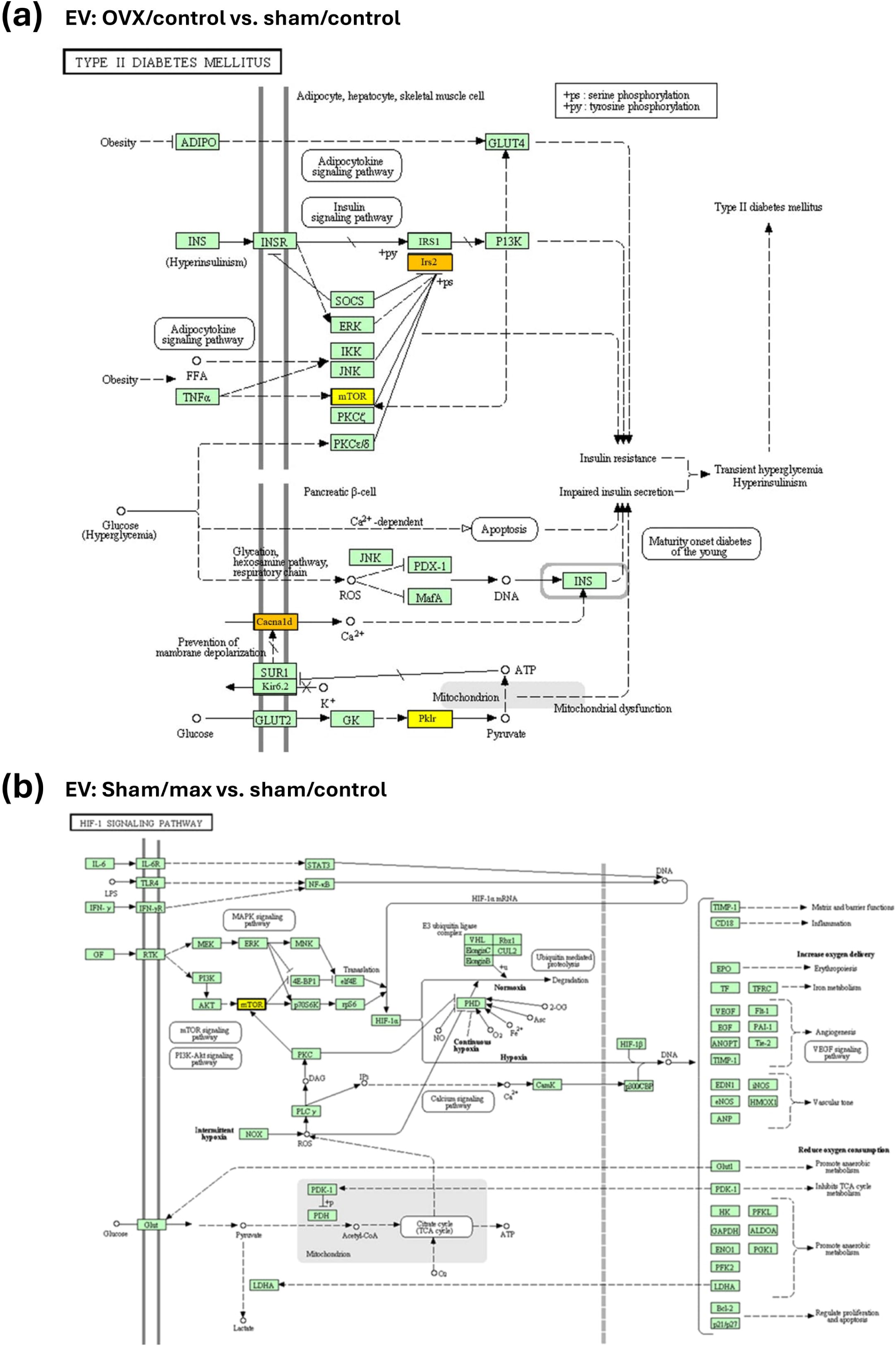
miR target signaling pathways (miRPath v3.0), illustrating pathways that union between EV miRs that were significantly different. (a) Visualization of type 2 diabetes signaling pathway and target proteins regulated by significantly different miRs carried by EV particles. (b) Visualization of HIF-1 signaling pathway and target proteins regulated by significantly different miRs carried by EV particles. Yellow=protein regulated by one miR, orange=protein regulated by more than one miR. OVX=ovariectomy, mTOR=mammalian target of rapamycin, IRS2=insulin receptor substrate 2, CACNA1D=calcium voltage-gated channel subunit alpha1 D, PKLR=pyruvate kinase L/R.

When studying the exercise-responsive EV miRs (sham/max vs. sham/control), miRPath v3.0 KEGG pathway analysis with pathways union option showed ten pathways, which were regulated by two of the miRs (Table S8). Of these, hypoxia inducible factor 1 (HIF-1) pathway, which plays a role in vascularization, is targeted by miR-100-5p, through its’ target protein mTOR (Figure 6b). When examining the two exercise-responsive HDL miRs, we observed 30 significant pathways, all of which were exclusively regulated by miR-100-5p, including type 2 diabetes pathway (Table S9).

To elucidate the possible functional relevance of the differentially expressed miRs in liver and muscle tissue, WB analysis was carried out for the most relevant target proteins from miRPath v3.0 analysis (Figure 6). Unfortunately, antibodies for CACNA1D proved to be defective, and therefore WB was carried out with mTOR (both muscle and liver), PKLR (only muscle), and IRS2 (only liver) (Figure 7a,b, Figure S4). The differentially expressed miRs and target protein expression in liver had no association (p≥0.126, Figure S4), but changes in the target proteins were observed in skeletal muscle tissue (Figure 7a,b,d,e). Muscle mTOR expression was lower in sham than in OVX (p=0.049, Figure 7a,e). There was an interaction of OVX and acute exercise in muscle PKLR expression (p=0.008, Figure 7b,e). mTOR is known for its diverse roles in metabolism regulation (Saxton & Sabatini, 2017), whereas PKLR takes part in energy production through glycolysis (Israelsen & Vander Heiden, 2015).

**FIGURE 7.**
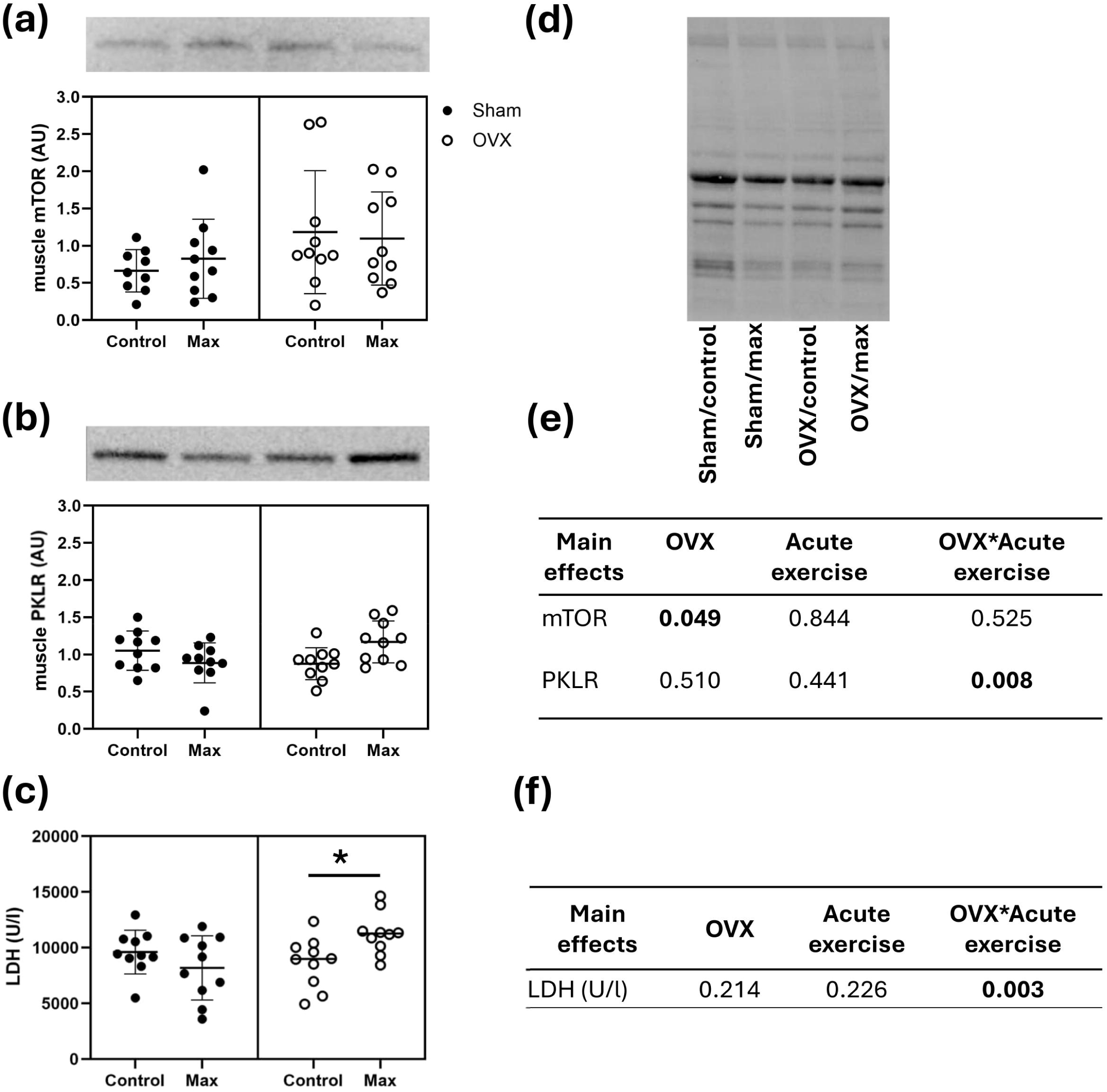
Loss of ovarian hormones and acute bout of exercise affect miR target protein levels in skeletal muscle. **(a)** WB analysis of mTOR level in muscle. **(b)** WB analysis of muscle PKLR. **(c)** Enzyme activity assay of LDH in muscle. **(d)** WB image of total protein, against which the results were normalized. **(e)** Main effects of OVX, acute exercise and their interaction on mTOR and PKLR levels in muscle (*p*-values). **(f)** Main effects of OVX, acute exercise and their interaction on LDH activity (p-values). OVX=ovariectomy, mTOR=mammalian target of rapamycin, PKLR=pyruvate kinase L/R, LDH=lactate dehydrogenase.

Since PKLR is an integral part of glycolysis, as it catalyzes the conversion of phosphoenolpyruvate (PEP) to pyruvate in the last step of glycolysis, we hypothesized that the change in PKLR induced by miR signaling changes would be reflected on the lactate dehydrogenase (LDH) enzyme activity, which is the next step in anaerobic glycolysis. LDH catalyzes the production of lactate from pyruvate, and its activity is a commonly used marker of the level of anaerobic energy production. The loss of ovarian hormones resulted in higher LDH activity in response to an acute bout of exercise (p=0.003) (Figure 7c,f).

## 4 DISCUSSION

In this study, we examined the effect of loss of ovarian hormones and acute exercise in the sRNA cargo of EV and HDL particles in female rats. We show for the first time that loss of ovarian hormones affects specifically the nucleic acid cargo of circulating EVs. We have demonstrated before that acute exercise-induced miR-response in EV and HDL particles is absent in postmenopausal women with low ovarian hormone levels (Karvinen et al., 2023), which was confirmed here in a rat model. We further demonstrate that the estrogen responsive miRs coordinates the glycolytic pathway in skeletal muscle. Finally, we show that the loss of ovarian hormones leads to higher anaerobic energy production during acute bout of exercise, suggesting an inferior ability to sustain aerobic energy production during exercise.

Loss of ovarian hormones increased body fat mass and reduced maximal running capacity as shown previously in our larger study cohort (E. Lee et al., 2024). Several studies have demonstrated that the loss of ovarian function in women and female rats is linked to weight gain, an increase in adipose tissue mass, and deterioration in metabolic health (Asarian & Geary, 2002; Juppi et al., 2022; E. Lee et al., 2024). Furthermore, loss of ovarian hormones has been associated with decreased physical activity levels in rodents (Chen et al., 2014; Park et al., 2016), and similar findings have been associated with menopause in women (Gabriel et al., 2015; Laakkonen et al., 2017). These findings together with our current results indicate inter-species similarity in responses to loss of ovarian hormones.

We show for the first time that loss of ovarian hormones changes specifically the nucleic acid cargo of EVs. Previous study by Sahu et al. observed that aging changed specifically the nucleic acid cargo of circulating EVs in a mouse model (Sahu et al., 2021). To our knowledge, we demonstrate for the first time that also loss of ovarian hormones leads to similar effects. Taken together, these findings indicate that aging-related alterations specifically impact the EV RNA cargo. Accordingly, we continued with detailed analysis of the sRNA species carried via EV and HDL particles.

In the present study the relative abundance of miRs was between 19-30% and it represented the first or second most abundant sRNA species of the top 100 host sRNAs. Further examination revealed that loss of ovarian hormones affected miRs associated with type 2 diabetes signaling pathways in circulating EVs. When comparing sham and OVX groups, we found nine miRs to be differently expressed in EVs, and 16 in HDL particles. Of the EV miRs, let-7g-5p, miR-100-5p, and miR-145-5p were associated to genes related to type 2 diabetes. In this pathway, let-7g-5p and miR-145-5p had targeted proteins CACNA1D and IRS2, while let-7g-5p targeted also PKLR. The target protein of miR-100-5p in this pathway was mTOR. Of these, let-7g-5p and miR-100-5p were also among the 20 most common miRs, highlighting their physiological relevance. In HDL particles, the differentially expressed miRs 100-5p and 99b-5p were associated with phosphoinositide 3-kinase (PI3K)/protein kinase B (PKB, otherwise known as AKT) signaling pathway. Both of the miRs had fibroblast growth factor receptor 3 (FGFR3), and miR-100-5p had also mTOR as a target protein. The PI3K/AKT pathway is mediated by insulin, affecting energy metabolism in insulin-sensitive tissues (Cao et al., 2023). In obesity, both up- and downregulation of the PI3K/AKT pathway have been found to be beneficial in a context-dependent manner (Savova et al., 2023). Our findings highlight the significant impact of loss of ovarian hormone on the nucleic acid cargo, particularly in miRs involved in type 2 diabetes and insulin signaling pathways, underscoring the complex interplay between hormonal changes and metabolic regulation.

Our previous research has shown that in postmenopausal women, low systemic estrogen levels reduce the miR-response to exercise in both EV and HDL particles (Karvinen et al., 2023). Our present results confirm that the loss of ovarian hormones blunts the exercise-induced miR response of EVs also in female rats. While in the sham groups there were three exercise-responsive miRs, none were observed in the OVX group. Interestingly, even though the OVX group had lower maximal running capacity in the POST test compared to sham, in the END measurement, where the running test was performed only once, there was no difference between the groups. This observation suggests that the difference between the systemic response to exercise in OVX rats is not attributable to a reduced running stimulus but rather represents a physiological change.

Of the exercise-responsive EV miRs in sham groups, miR-100-5p was upregulated and found to be associated with hypoxia-inducible factor-1 (HIF-1) signaling through mTOR. However, we did not observe a difference in the mTOR level in skeletal muscle in response to exercise. HIF-1 target genes act in skeletal muscle in oxygen transport enhancement through e.g., activation of angiogenesis and glycolysis (Lindholm & Rundqvist, 2015). The upregulation of miRs targeting HIF-1 in sham/max group may indicate attenuated HIF-1 response to acute exercise stimulus.

Excitingly, there was an interaction of OVX and acute exercise on the level of PKLR in skeletal muscle. It seems that in response to exercise, the PKLR level in muscle decreases in sham group, whereas in OVX the PKLR level increases. However, in addition to the main effect of OVX and acute exercise interaction, no significant findings between the control and corresponding max groups were observed. PKLR catalyzes the second last step of anaerobic glycolysis, which converts phosphoenolpyruvate (PEP) to pyruvate in ATP production. Hence, elevated PKLR levels may indicate increased glycolytic activity, likely as a response to higher energy demands in the muscle. LDH in turn catalyzes the last step of anaerobic glycolysis, converting pyruvate to lactate. In our study, the OVX group exhibited higher LDH activity in the max group, further indicating that the glycolytic flux is higher in response to acute exercise with low ovarian hormone status. Of the ovarian hormones, 17β-estradiol (E2) has been proven to act as key regulator of energy homeostasis affecting several tissues, including skeletal muscle (Mauvais-Jarvis et al., 2013). E2 promotes lipid oxidation in skeletal muscle in vivo and in vitro (Garrido et al., 2014; Hamadeh et al., 2005), while OVX may lead to reduced oxidative capacity due to impaired mitochondrial function (Hu et al., 2024; M. J. Torres et al., 2018). Our results are in line with previous literature, yet to our knowledge, our study is the first to show that loss of ovarian hormones leads to higher anaerobic energy production in response to acute bout of exercise. Our results suggest that OVX rats are less effective at sustaining aerobic energy production during an acute bout of exercise.

We observed one exercise-responsive miR in the HDL particles in the OVX group, while there were two such miRs in the sham group. In the OVX group, miR-200b-3p was downregulated in response to exercise (OVX/max vs. OVX/control). Patients with type 2 diabetes have been shown to have less circulating miR-200b in plasma than healthy controls (Dantas da Costa e Silva et al., 2019). Also, coronary artery disease patients have been shown to display downregulation of miR-200b-3p after acute bout of exercise (Mayr et al., 2019). In cultured alveolar cells, miR-200b-3p has been shown to be capable of decreasing senescence markers and restoring regenerative potential (Moimas et al., 2019). Taken together, these findings suggest that the downregulation of miR-200b-3p may be linked to negative cardiovascular factors. Our finding of the downregulation of this specific miR in HDL particles in OVX group may indicate that miR-200b-3p is exercise-responsive and is affected by the loss of ovarian hormones, possibly providing a link between unfavorable cardiovascular changes associated with the loss of ovarian hormones.

In EV and HDL particles, the relative abundance of rDR was between 49-57% and it represented the first or second most abundant sRNA species of the top 100 host sRNAs. Interestingly, we found significant differences in rDR species only in the HDL particles when comparing the OVX groups (max vs. control), suggesting that the loss of ovarian hormones together with acute exercise modulates the rDR cargo of HDL, whereas acute exercise, or OVX alone do not cause changes. All of these rDRs originated from SSU rRNA. SSU rRNA is one of the two main RNA components of a ribosome, also known as 18S rRNA. SSU rRNA plays a crucial role in the translation of genetic information into proteins. In our previous study the majority of the exercise-responsive rDR species in HDL particles belonged to mitochondrial ribosomal subunits (Karvinen et al., 2023). While mitochondrial peptides are now known to regulate metabolism (Kim et al., 2017), the potential role of SSU rRNA fragments in signaling remains uncovered.

In EV and HDL particles, the relative abundance of tDR was between 6-14% and it represented the third most abundant sRNA species of the top 100 host sRNAs. In EVs, tDR derived from tRNA-Met-CAT-3-1 was higher both in OVX rats compared with sham and in sham/max compared with sham/control group. In addition, tDR derived from tRNA-Met-CAT-1-1 was upregulated in OVX rats compared with sham. However, no effect of acute exercise in EV tDRs was seen when comparing OVX/max to OVX/control, tRNA fragments, while still under thorough investigation, have been found to play significant roles in systemic signaling, impacting various biological processes (Kaimwal et al., 2025; Torres & Martí, 2021). Their currently known biological roles include regulating gene expression by inhibition of translation (Lee et al., 2009; Li et al., 2012). Although tDRs may have similar regulatory functions to miRs, their role as systemic signaling molecules remains unclear.

To conclude, this study highlights the role of ovarian hormones in regulating miR signaling in response to acute exercise. Accordingly, the miR cargo of circulating EVs may affect the protein expression of the target tissues affecting metabolism. Our findings emphasize the importance of ovarian hormones in the acute response to exercise and their potential role in facilitating the health benefits associated with exercise.

## STRENGTHS AND LIMITATIONS

The present study was designed to investigate the effects of OVX and acute bout of exercise on systemic signaling. The first key strength of the current study is that the animals were fully grown adults (∼7 months old) before undergoing surgery, ensuring a more robust model of menopause. Second, to ensure a robust difference in the ovarian hormone levels between OVX and sham groups, the rats in the sham group were euthanized at the time of proestrus. The third strength of the study is the implementation of allocating animals into the four study groups matched for body weight and maximal running capacity. Fourth, we utilized an isolation protocol validated by us enabling the separation of EV and HDL fractions before RNA isolation. Our study also has certain limitations. Our study concentrates only on female sex, which was necessary for our study question. Due to the low protein content of the EV samples, we were not able to carry out the protein analysis via WB and hence utilized DB analysis when necessary. Nevertheless, total protein, DB, and EM analysis confirmed that the EV and HDL particles were enriched in separate fractions. We were not able to detect all the target proteins in both muscle and liver tissues due to defectiveness or tissue specificity of the antibodies. Also, due to the limited sample volume we were not able to assess the number of EV or HDL particles in the samples. Nevertheless, we are confident that the results are relevant regardless of whether the observed differences between the groups are partly due to the increased number of particles, as only a few specific miRs showed a clear response to exercise stimulus in sham group, supporting highly coordinated regulatory mechanisms for miR packaging. Likely due to high lipid and protein content, we were not able to perform Raman spectroscopy for HDL particles. Also, we were not able to trace the EV and HDL particles to determine their target tissues, yet based on previous literature, we are confident that liver and muscle are affected by systemic signaling.

## DATA AVAILABILITY STATEMENT

The results obtained are presented in the manuscript. When applicable, a link for the data source is presented.

## FUNDING STATEMENT

This study was funded by grants from the Research Council of Finland (grant numbers #332946 and #354603 to SK and grant number #341058 to ML).

## ACKNOWLEDGEMENTS

We would like to thank the laboratory staff at the Faculty of Sport and Health Sciences for their invaluable assistance in the data collection. We acknowledge the services of university of Helsinki: EV Core in FIMM Technology Centre supported by HiLIFE and Biocentre Finland for performing electron microscopy work and Electron Microscopy Unit of the Institute of Biotechnology for providing the facilities.

## DECLARATION OF INTEREST STATEMENT

The authors declare no conflict of interest.

## AUTHOR CONTRIBUTIONS

VP: Methodology (WB, miR-pathway); analysis; writing - original draft. AM: Methodology (EV analysis); analysis; writing (supportive). T-MK: Analysis; writing (review and editing). TAN: methodology (animal study); writing (review and editing). EH: Methodology (Raman spectroscopy); analysis; writing (review and editing). JI: Resources (Raman spectroscopy); methodology; writing (review and editing). ML: Methodology; supervision (supportive); writing (review and editing). SK: Conceptualization (lead); resources (lead); supervision (lead); writing - review and editing (lead).

## Notes

### Competing Interest Statement

The authors have declared no competing interest.

