## Supplemental figures and tables for "Loss of ovarian hormones modulates the nucleic acid content of circulating extracellular vesicles and skeletal muscle metabolism in response to acute exercise"

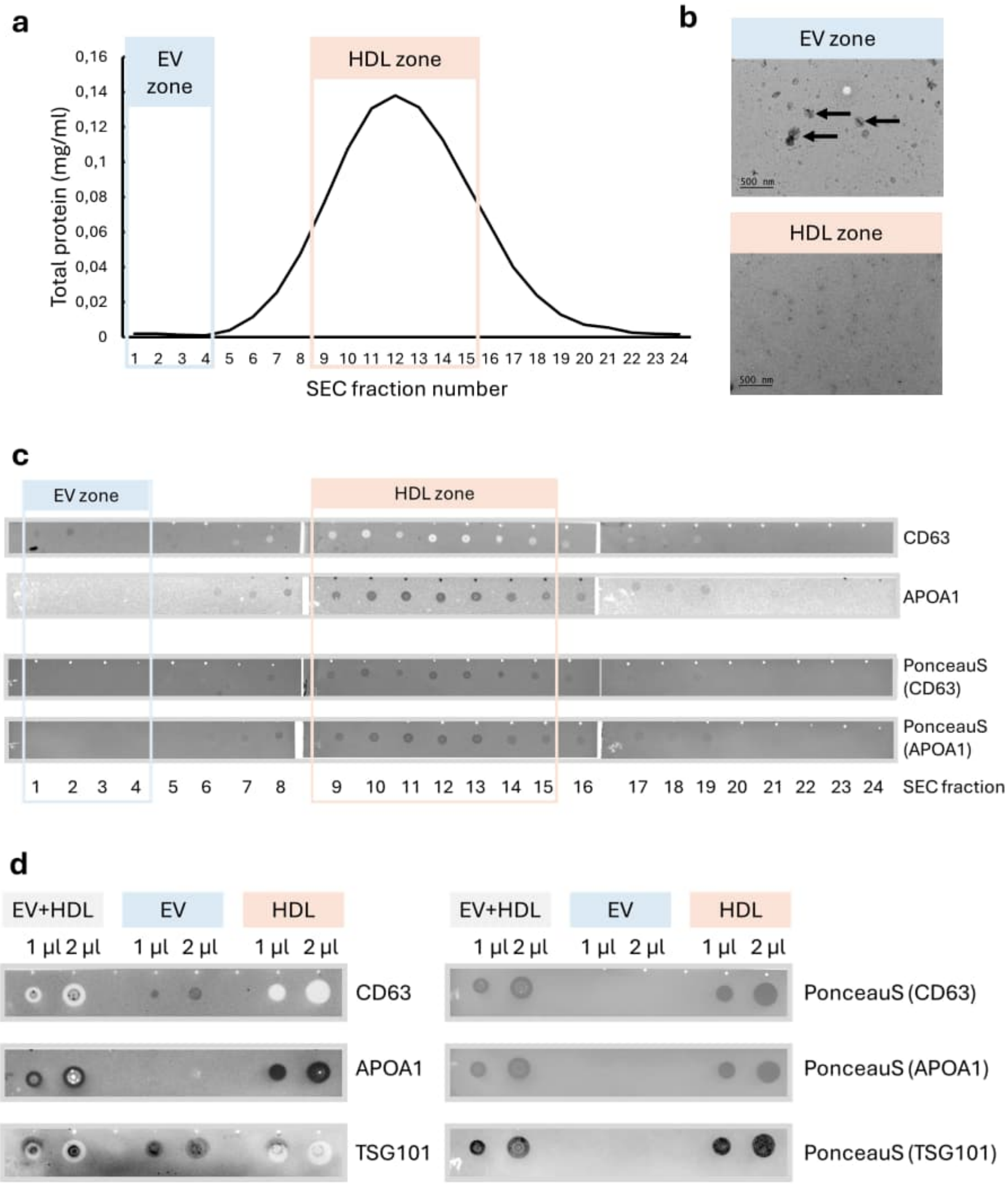


**FIGURE S1. EV and HDL particles enrich in separate fractions. (a)** Total protein (BCA) analysis of SEC fractions 1–24 from a representative EV+HDL fraction from plasma. **(b)** Representative EM images of EV and HDL fractions isolated from plasma. **(c)** DB verification of enrichment of EVs into SEC fractions 1–4 with CD63 antibody and HDL particles into SEC fractions 9–15 with APOA1 antibody with corresponding total protein blots stained with PonceauS from a representative EV and HDL fraction isolated from plasma. **(e)** Fractions EV+HDL, EV and HDL separated via SEC and visualized with 1 and 2 µL sample with CD63, APOA1, and TSG101 antibodies (left panel) and corresponding total protein blots stained with PonceauS (right panel) from a representative EV and HDL fractions isolated from plasma. EV=extracellular vesicle, HDL=high-density lipoprotein, SEC=size-exclusion chromatography.

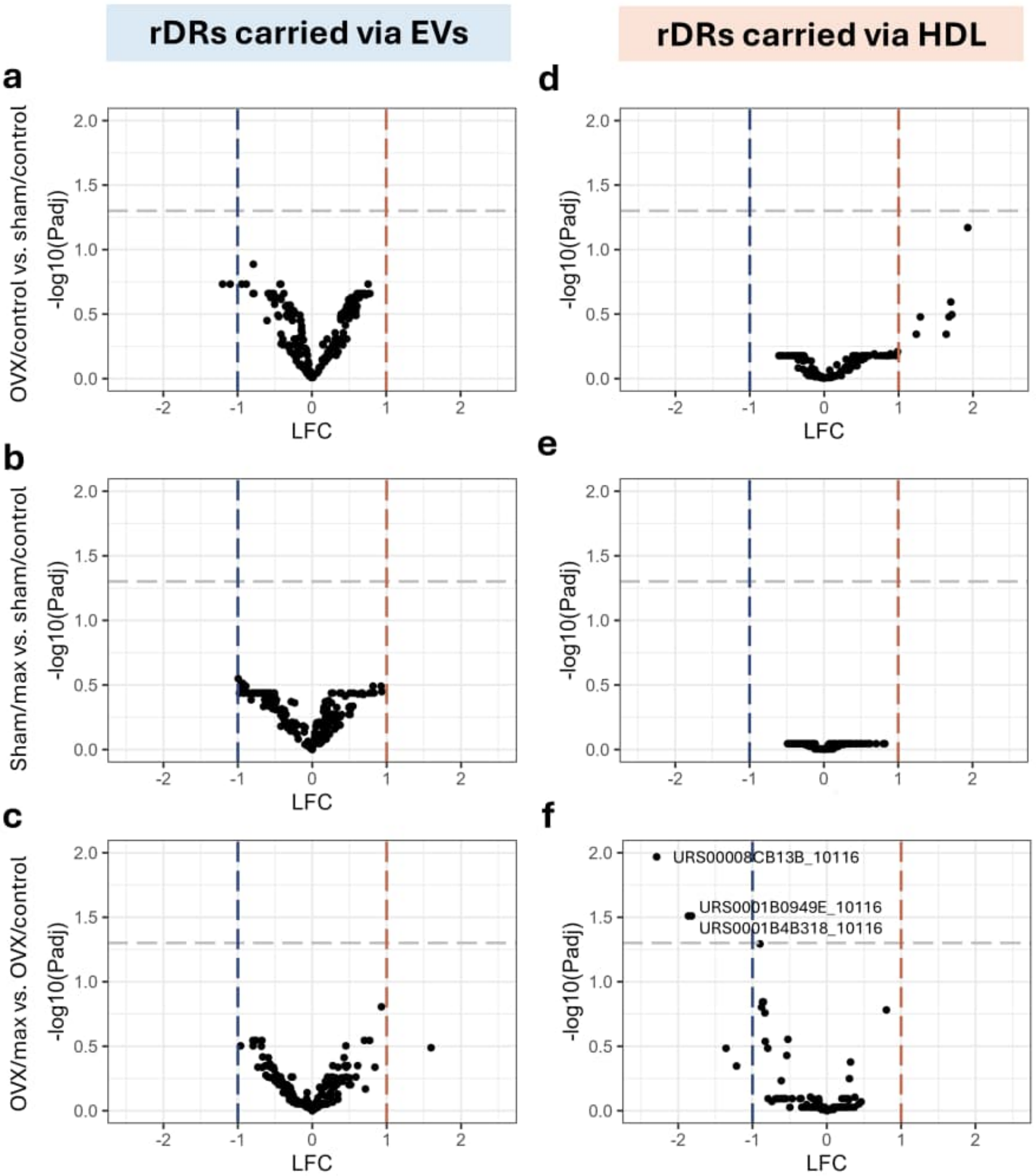


**FIGURE S2**. **The rDR cargo of EV and HDL particles responds to acute bout of exercise differently. (a-c)** Volcano plots showing group comparisons of differential expression of rDRs in EV particles. **(d-f)** Volcano plots showing group comparisons of differential expression of rDRs in HDL particles. OVX=ovariectomy, rDR=ribosomal RNA (rRNA) -derived sRNA, LFC=log fold-change.

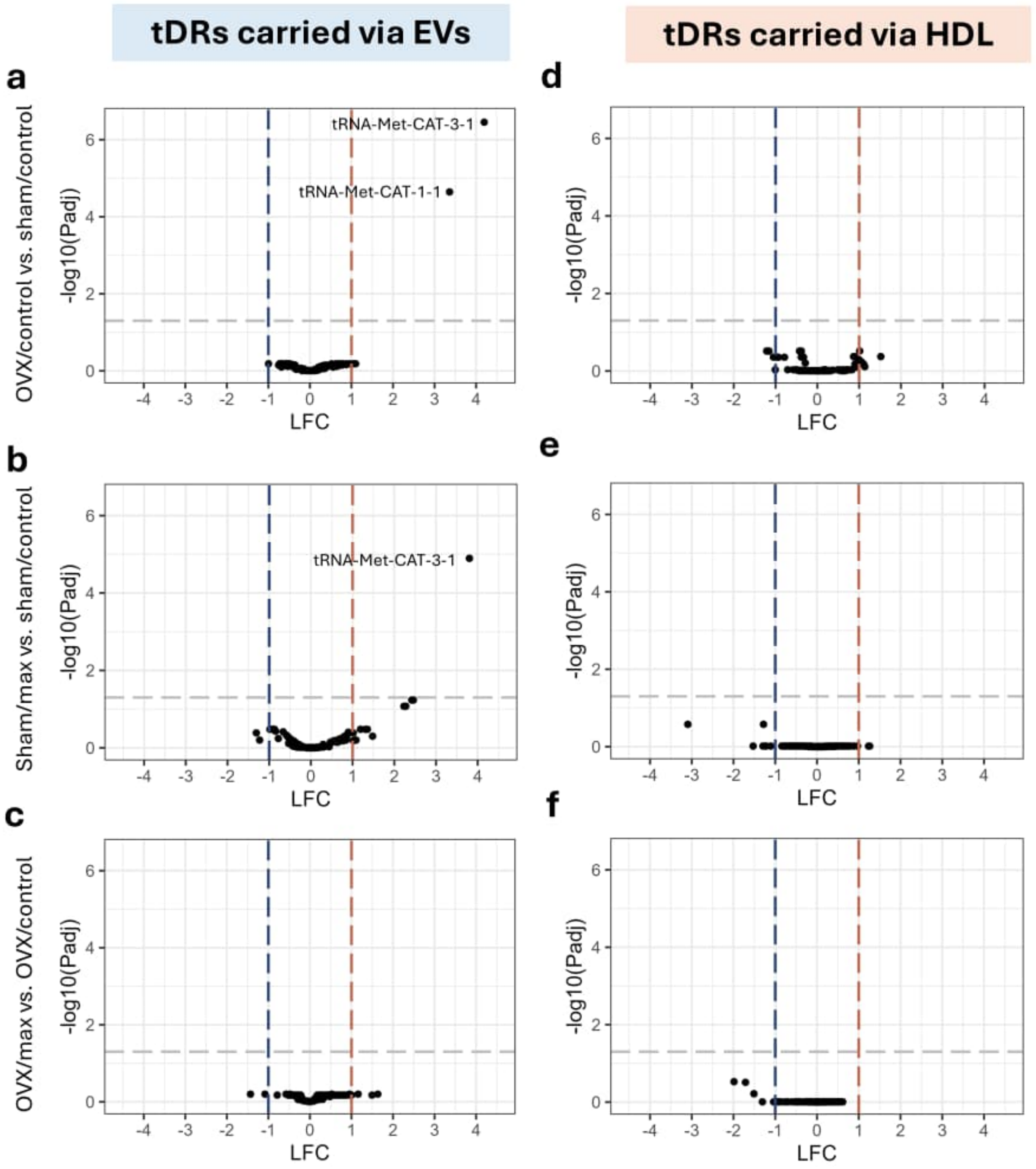


**FIGURE S3**. **The tDR cargo of EV and HDL particles responds to acute bout of exercise differently. (a-c)** Volcano plots showing group comparisons of differential expression of tDRs in EV particles. **(d-f)** Volcano plots showing group comparisons of differential expression of tDRs in HDL particles. OVX=ovariectomy, tDR=transfer RNA (tRNA) -derived sRNA, LFC=log fold-change.

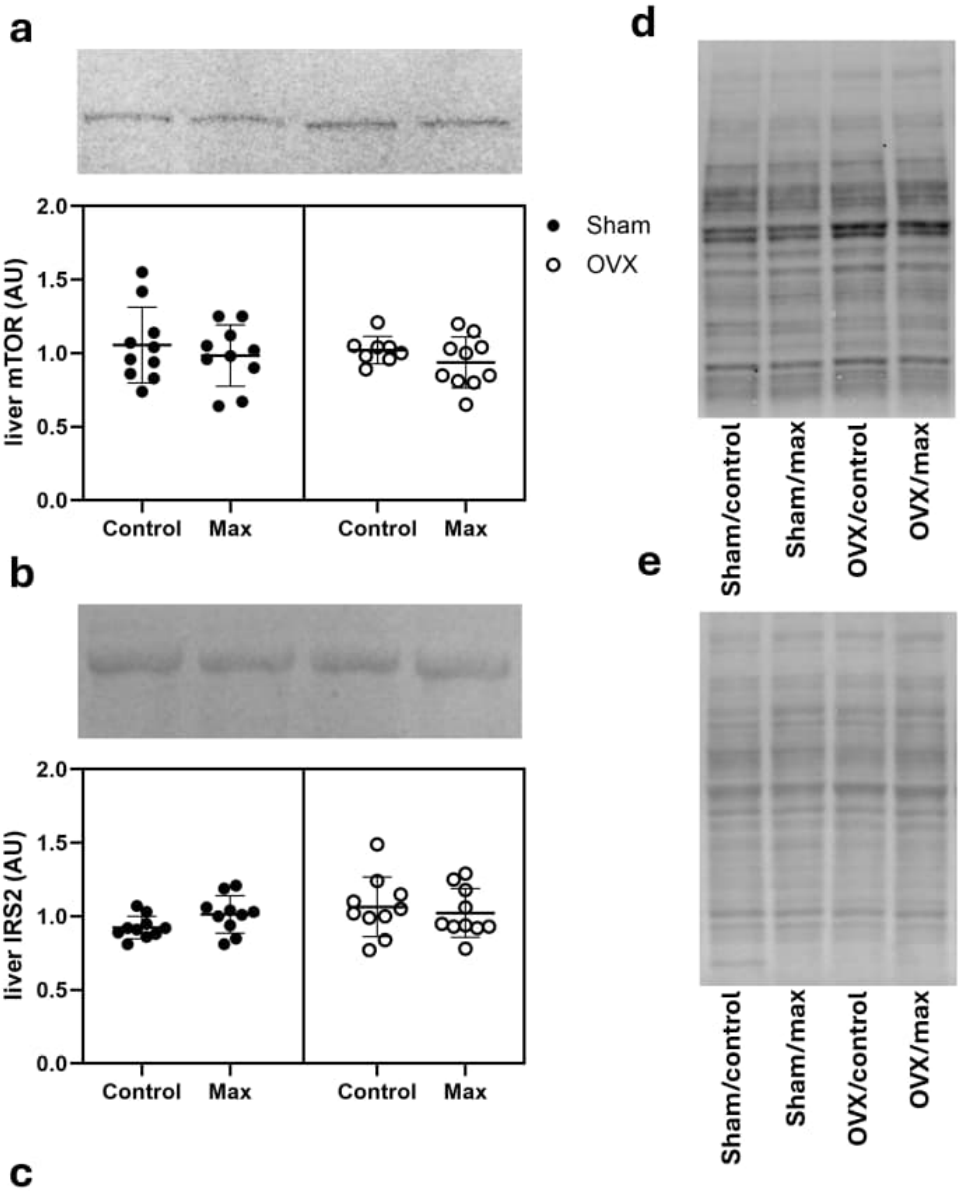


c

| Main effects | OVX | Acute exercise | OVX*Acute exercise |
|---|---|---|---|
| mTOR | 0.544 | 0.239 | 0.942 |
| IRS2 | 0.126 | 0.629 | 0.178 |

**FIGURE S4**. **Loss of ovarian hormones and acute bout of exercise did not affect the miR target protein levels in liver.** **(a)** WB analysis of mTOR level in liver. **(b)** WB analysis of muscle IRS2 level in liver. **(c)** Main effects of OVX, acute exercise and their interaction on mTOR and IRS2 levels in liver. **(d-e)** Corresponding WB images of total protein, against which the results were normalized. OVX=ovariectomy, mTOR=mammalian target of rapamycin, IRS2= insulin receptor substrate 2.

### TABLES

**Table S1.** miR counts in EV and HDL particles in the study groups.

| | | EV | | | | HDL | | | |
|---|---|---|---|---|---|---|---|---|---|
| Number | miR | Sham/control | Sham/max | OVX/control | OVX/max | Sham/control | Sham/max | OVX/control | OVX/max |
| 1 | let-7a-5p | 33610 | 54906 | 58282 | 57313 | 24073 | 27751 | 29935 | 41685 |
| 2 | let-7b-5p | 45021 | 59732 | 64835 | 52030 | 31015 | 43941 | 39991 | 62414 |
| 3 | let-7c-2-3p | 71 | 145 | 182 | 190 | | | | |
| 4 | let-7c-5p | 25250 | 43464 | 40962 | 36856 | 21470 | 30590 | 25387 | 39516 |
| 5 | let-7d-3p | 6476 | 10640 | 10935 | 9486 | 8172 | 11896 | 9115 | 13803 |
| 6 | let-7d-5p | 6583 | 10873 | 11244 | 10010 | 7876 | 8678 | 7439 | 9487 |
| 7 | let-7e-5p | 1245 | 1898 | 1692 | 1708 | 831 | 849 | 862 | 1201 |
| 8 | let-7f-5p | 49439 | 77149 | 77945 | 77001 | 27636 | 35066 | 38822 | 60539 |
| 9 | let-7g-5p | 9834 | 10823 | 10587 | 15947 | 7211 | 6879 | 5349 | 4563 |
| 10 | let-7i-5p | 110030 | 152200 | 170430 | 145955 | 94198 | 114086 | 95914 | 139715 |
| 11 | miR-100-5p | 1823 | 14890 | 8578 | 12454 | 5731 | 3427 | 793 | 564 |
| 12 | miR-103-3p | 773 | 1138 | 1098 | 1449 | 993 | 800 | 607 | 740 |
| 13 | miR-101a-3p | 256 | 357 | 554 | 596 | | | | |
| 14 | miR-106b-3p | 276 | 540 | 511 | 449 | 217 | 397 | 339 | 312 |
| 15 | miR-101b-3p | 73 | 87 | 148 | 111 | | | | |
| 16 | miR-10a-5p | 18255 | 30909 | 23445 | 31622 | 4308 | 4443 | 5412 | 8885 |
| 17 | miR-122-5p | 24484 | 67542 | 56950 | 20219 | 46599 | 26837 | 38543 | 16884 |
| 18 | miR-125a-5p | 3486 | 5752 | 4836 | 7268 | 1584 | 1313 | 987 | 1904 |
| 19 | miR-125b-5p | 448 | 872 | 636 | 943 | 1618 | 1037 | 261 | 450 |
| 20 | miR-126a-3p | 20634 | 28338 | 24071 | 31638 | 4711 | 6256 | 5407 | 7744 |

| | | | | | | | | | |
|---|---|---|---|---|---|---|---|---|---|
| 21 | miR-127-3p | 113 | 282 | 420 | 534 | 404 | 290 | 145 | 185 |
| 22 | miR-128-3p | 51775 | 82806 | 85128 | 71748 | 88676 | 114676 | 86417 | 108088 |
| 23 | miR-139-5p | 624 | 1229 | 852 | 846 | 211 | 317 | 356 | 441 |
| 24 | miR-1-3p | 845 | 1352 | 1280 | 1421 | 1088 | 644 | 598 | 2162 |
| 25 | miR-132-3p | 114 | 144 | 258 | 279 | | | | |
| 26 | miR-142-3p | 596 | 583 | 816 | 1183 | 317 | 414 | 288 | 462 |
| 27 | miR-133a-3p | 299 | 198 | 56 | 341 | | | | |
| 28 | miR-143-3p | 27269 | 26926 | 36466 | 20871 | 16519 | 10529 | 10151 | 6960 |
| 29 | miR-146a-5p | 1786 | 1601 | 1627 | 2593 | 486 | 718 | 600 | 773 |
| 30 | miR-146b-5p | 2727 | 4258 | 6622 | 15744 | 1075 | 1218 | 1687 | 2374 |
| 31 | miR-141-3p | 120 | 101 | 134 | 320 | | | | |
| 32 | miR-148a-3p | 754 | 1117 | 1233 | 821 | 300 | 369 | 329 | 287 |
| 33 | miR-148a-5p | 190 | 205 | 253 | 295 | 171 | 202 | 262 | 257 |
| 34 | miR-142-5p | 98 | 254 | 408 | 362 | | | | |
| 35 | miR-148b-3p | 231318 | 364421 | 413206 | 330739 | 220010 | 296548 | 265971 | 314655 |
| 36 | miR-150-5p | 803 | 1342 | 1403 | 2412 | 554 | 714 | 570 | 303 |
| 37 | miR-145-5p | 327 | 197 | 123 | 37 | | | | |
| 38 | miR-151-3p | 6648 | 11039 | 10861 | 9554 | 7764 | 10150 | 8099 | 9850 |
| 39 | miR-152-3p | 142 | 161 | 208 | 216 | 174 | 116 | 76 | 79 |
| 40 | miR-155-5p | 246 | 870 | 1615 | 4677 | 185 | 202 | 207 | 40 |
| 41 | miR-16-5p | 427 | 378 | 398 | 593 | 192 | 177 | 147 | 157 |
| 42 | miR-17-5p | 352 | 585 | 580 | 773 | 460 | 383 | 506 | 349 |
| 43 | miR-181a-5p | 824 | 1473 | 1427 | 2241 | 1331 | 1461 | 927 | 1110 |
| 44 | miR-181b-5p | 291 | 504 | 432 | 414 | 278 | 295 | 270 | 457 |
| 45 | miR-181d-5p | 466 | 891 | 707 | 670 | 952 | 827 | 967 | 1075 |
| 46 | miR-182 | 89 | 367 | 325 | 471 | 339 | 244 | 36 | 138 |
| 47 | miR-184 | 3771 | 8001 | 5312 | 5505 | 1653 | 8846 | 1797 | 8447 |
| 48 | miR-185-5p | 248 | 335 | 332 | 490 | 443 | 335 | 383 | 434 |
| 49 | miR-186-5p | 522 | 877 | 1235 | 1353 | 336 | 485 | 458 | 677 |

| | | | | | | | | | |
|---|---|---|---|---|---|---|---|---|---|
| 50 | miR-191a-3p | 317 | 399 | 544 | 360 | 562 | 789 | 602 | 792 |
| 51 | miR-191a-5p | 25681 | 39373 | 39575 | 40445 | 49910 | 50594 | 50778 | 52992 |
| 52 | miR-192-5p | 212 | 355 | 438 | 334 | 293 | 201 | 1203 | 127 |
| 53 | miR-199a-3p | 583 | 497 | 486 | 438 | 1199 | 1017 | 200 | 164 |
| 54 | miR-183-5p | 87 | 231 | 451 | 720 | | | | |
| 55 | miR-199a-5p | 190 | 302 | 222 | 636 | 369 | 264 | 98 | 44 |
| 56 | miR-1b | 22681 | 10881 | 4690 | 10326 | 4739 | 2267 | 3014 | 3098 |
| 57 | miR-200b-3p | 628 | 827 | 1528 | 1995 | 2535 | 1376 | 528 | 72 |
| 58 | miR-206-3p | 200 | 1107 | 1048 | 1385 | 1093 | 574 | 355 | 1635 |
| 59 | miR-20a-5p | 481 | 576 | 824 | 1061 | 417 | 450 | 469 | 471 |
| 60 | miR-210-3p | 205 | 243 | 294 | 411 | 308 | 299 | 184 | 213 |
| 61 | miR-21-5p | 32896 | 55624 | 69432 | 101693 | 31153 | 32026 | 21897 | 27841 |
| 62 | miR-221-3p | 221 | 735 | 908 | 1700 | 757 | 600 | 320 | 251 |
| 63 | miR-223-5p | 2424 | 3092 | 3539 | 4661 | 1243 | 1580 | 1484 | 2500 |
| 64 | miR-22-3p | 1107 | 1876 | 1873 | 1994 | 996 | 1103 | 1509 | 1268 |
| 65 | miR-200a-3p | 271 | 276 | 241 | 263 | | | | |
| 66 | miR-22-5p | 153 | 358 | 474 | 198 | 164 | 172 | 189 | 215 |
| 67 | miR-23a-3p | 172 | 107 | 214 | 248 | 294 | 250 | 115 | 96 |
| 68 | miR-203a-3p | 85 | 407 | 213 | 230 | | | | |
| 69 | miR-24-2-5p | 336 | 416 | 406 | 403 | 341 | 364 | 361 | 357 |
| 70 | miR-204-3p | 84 | 72 | 308 | 177 | | | | |
| 71 | miR-24-3p | 4838 | 6906 | 7251 | 7574 | 4868 | 5700 | 4765 | 5785 |
| 72 | miR-25-3p | 2618 | 3845 | 4519 | 5839 | 2246 | 2751 | 2503 | 2643 |
| 73 | miR-26a-5p | 67717 | 120248 | 120772 | 123020 | 36106 | 46237 | 49017 | 67410 |
| 74 | miR-26b-5p | 393 | 631 | 725 | 744 | 319 | 348 | 315 | 279 |
| 75 | miR-27a-3p | 2330 | 4123 | 5608 | 7006 | 2547 | 2942 | 2140 | 1967 |
| 76 | miR-218a-5p | 79 | 286 | 244 | 69 | | | | |
| 77 | miR-27a-5p | 219 | 361 | 455 | 595 | 148 | 169 | 195 | 276 |
| 78 | miR-27b-3p | 2175 | 2217 | 2563 | 2402 | 4463 | 2696 | 1144 | 892 |

| | | | | | | | | | |
|---|---|---|---|---|---|---|---|---|---|
| 79 | miR-28-3p | 419 | 686 | 991 | 1197 | 617 | 568 | 642 | 658 |
| 80 | miR-29a-3p | 357 | 372 | 482 | 693 | 377 | 301 | 215 | 167 |
| 81 | miR-30a-3p | 252 | 482 | 570 | 446 | 260 | 232 | 193 | 224 |
| 82 | miR-30a-5p | 3164 | 3078 | 3763 | 3083 | 1499 | 931 | 817 | 1078 |
| 83 | miR-30c-5p | 3279 | 3642 | 7399 | 16656 | 2197 | 2280 | 2563 | 2543 |
| 84 | miR-30d-5p | 11005 | 16220 | 18384 | 19473 | 5986 | 5915 | 6505 | 7952 |
| 85 | miR-30e-3p | 562 | 1087 | 2771 | 5732 | 393 | 458 | 586 | 638 |
| 86 | miR-30e-5p | 1400 | 2365 | 4135 | 7303 | 447 | 546 | 836 | 995 |
| 87 | miR-320-3p | 5480 | 7926 | 8065 | 7961 | 6219 | 6209 | 7617 | 6194 |
| 88 | miR-328a-3p | 8198 | 12513 | 11703 | 10188 | 20721 | 19495 | 17312 | 19409 |
| 89 | miR-330-3p | 815 | 1246 | 1312 | 1002 | 1025 | 1109 | 1028 | 1550 |
| 90 | miR-28-5p | 70 | 126 | 195 | 150 | | | | |
| 91 | miR-339-3p | 120 | 125 | 259 | 314 | 201 | 274 | 228 | 308 |
| 92 | miR-339-5p | 104 | 284 | 263 | 422 | 506 | 495 | 365 | 512 |
| 93 | miR-340-3p | 550 | 852 | 1066 | 1305 | 269 | 315 | 251 | 418 |
| 94 | miR-340-5p | 429 | 501 | 790 | 1121 | 192 | 168 | 109 | 284 |
| 95 | miR-342-5p | 1231 | 1852 | 2135 | 1660 | 977 | 1336 | 914 | 1981 |
| 96 | miR-31a-5p | 31 | 114 | 148 | 142 | | | | |
| 97 | miR-361-3p | 562 | 730 | 866 | 890 | 618 | 534 | 462 | 1018 |
| 98 | miR-378a-3p | 997 | 514 | 341 | 362 | 330 | 234 | 235 | 210 |
| 99 | miR-423-3p | 5051 | 8541 | 10418 | 13375 | 9351 | 10614 | 9454 | 10269 |
| 100 | miR-423-5p | 28116 | 39544 | 40552 | 31661 | 28736 | 36733 | 32985 | 43251 |
| 101 | miR-425-5p | 3405 | 5806 | 4869 | 4932 | 7046 | 6651 | 5938 | 8300 |
| 102 | miR-451-5p | 11478 | 18184 | 10075 | 6031 | 1985 | 2730 | 4908 | 4330 |
| 103 | miR-484 | 251 | 562 | 255 | 909 | 577 | 559 | 409 | 719 |
| 104 | miR-486 | 20731 | 36551 | 25560 | 27950 | 9529 | 18489 | 13482 | 20936 |
| 105 | miR-34c-5p | 381 | 878 | 1141 | 509 | | | | |
| 106 | miR-3559-3p | 153 | 200 | 145 | 156 | | | | |
| 107 | miR-375-3p | 328 | 976 | 1136 | 703 | | | | |

| | | | | | | | | | |
|---|---|---|---|---|---|---|---|---|---|
| 108 | miR-488-3p | 50 | 232 | 214 | 195 | | | | |
| 109 | miR-499-5p | 744 | 1403 | 1983 | 1399 | 219 | 109 | 267 | 162 |
| 110 | miR-532-5p | 223 | 330 | 463 | 774 | 270 | 218 | 183 | 150 |
| 111 | miR-674-3p | 466 | 623 | 596 | 388 | 421 | 570 | 445 | 396 |
| 112 | miR-676 | 217 | 181 | 179 | 178 | | | | |
| 113 | miR-7a-5p | 4656 | 6169 | 6368 | 12890 | 2589 | 3465 | 2689 | 4636 |
| 114 | miR-872-5p | 182 | 319 | 295 | 343 | 238 | 242 | 306 | 408 |
| 115 | miR-92a-3p | 140 | 263 | 432 | 444 | 123 | 155 | 246 | 280 |
| 116 | miR-92b-3p | 1594 | 3159 | 4246 | 6237 | 1291 | 1603 | 1919 | 1733 |
| 117 | miR-93-5p | 227 | 575 | 307 | 401 | 248 | 335 | 157 | 282 |
| 118 | miR-98-5p | 1299 | 2248 | 2461 | 2225 | 1404 | 1838 | 1688 | 2233 |
| 119 | miR-99a-5p | 17488 | 30467 | 32961 | 22403 | 7343 | 6563 | 6126 | 5264 |
| 120 | miR-99b-5p | 2020 | 3320 | 4289 | 6080 | 1305 | 1212 | 929 | 1076 |
| 121 | miR-9a-5p | 944 | 1849 | 1833 | 2326 | 525 | 507 | 236 | 401 |

**Table S2.** Twenty most significantly different miRs carried via EVs.

| EV | Number | miR | log2 Fold change | SE (log Fold change) | Adjusted p-value |
|---|---|---|---|---|---|
| OVX/control | **1** | **miR-1b** | **-2.958** | **0.371** | **0.000** |
| vs. | **2** | **miR-378a-3p** | **-2.259** | **0.402** | **0.000** |
| Sham/control | **3** | **miR-133a-3p** | **-3.333** | **0.595** | **0.000** |
| | **4** | **miR-100-5p** | **1.715** | **0.335** | **0.000** |
| | **5** | **miR-145-5p** | **-2.574** | **0.579** | **0.000** |
| | **6** | **let-7g-5p** | **-0.558** | **0.146** | **0.003** |
| | **7** | **miR-206-3p** | **2.237** | **0.668** | **0.014** |
| | **8** | **miR-30c-5p** | **-0.538** | **0.178** | **0.037** |
| | **9** | **miR-146a-5p** | **-0.900** | **0.302** | **0.039** |
| | 10 | miR-221-3p | 1.472 | 0.538 | 0.075 |
| | 11 | miR-155-5p | 1.618 | 0.686 | 0.170 |
| | 12 | miR-182 | 1.656 | 0.691 | 0.170 |
| | 13 | miR-92b-3p | 0.648 | 0.274 | 0.170 |
| | 14 | miR-183-5p | 1.852 | 0.810 | 0.191 |
| | 15 | miR-31a-5p | 2.079 | 0.941 | 0.220 |
| | 16 | miR-22-5p | 1.789 | 0.820 | 0.221 |
| | 17 | miR-375-3p | 1.282 | 0.627 | 0.274 |
| | 18 | miR-92a-3p | 1.036 | 0.503 | 0.274 |
| | 19 | miR-16-5p | -0.810 | 0.415 | 0.326 |
| | 20 | miR-30a-3p | 0.810 | 0.432 | 0.369 |
| Sham/max | **1** | **miR-100-5p** | **2.873** | **0.556** | **0.000** |
| vs. | **2** | **miR-221-3p** | **1.782** | **0.526** | **0.035** |
| Sham/control | **3** | **miR-127-3p** | **2.813** | **0.845** | **0.035** |
| | 4 | miR-1b | -1.265 | 0.425 | 0.086 |
| | 5 | miR-30c-5p | -0.544 | 0.187 | 0.086 |
| | 6 | miR-203a-3p | 1.947 | 0.758 | 0.162 |
| | 7 | miR-92b-3p | 0.637 | 0.250 | 0.162 |
| | 8 | miR-31a-5p | 3.005 | 1.175 | 0.162 |
| | 9 | miR-378a-3p | -1.111 | 0.450 | 0.182 |
| | 10 | let-7g-5p | -0.388 | 0.163 | 0.189 |
| | 11 | miR-375-3p | 1.482 | 0.621 | 0.189 |
| | 12 | miR-182 | 1.615 | 0.787 | 0.349 |
| | 13 | miR-206-3p | 1.218 | 0.594 | 0.349 |
| | 14 | miR-7a-5p | -0.422 | 0.201 | 0.349 |
| | 15 | miR-142-3p | -0.663 | 0.349 | 0.385 |
| | 16 | miR-143-3p | -0.418 | 0.212 | 0.385 |
| | 17 | miR-183-5p | 1.662 | 0.869 | 0.385 |
| | 18 | miR-223-5p | -0.517 | 0.266 | 0.385 |
| | 19 | miR-155-5p | 0.891 | 0.491 | 0.415 |
| | 20 | miR-27a-3p | 0.360 | 0.196 | 0.415 |
| OVX/max | 1 | miR-122-5p | -1.398 | 0.475 | 0.395 |
| vs. | 2 | miR-30e-3p | 1.246 | 0.470 | 0.489 |
| OVX/control | 3 | miR-484 | 1.284 | 0.530 | 0.622 |

| | | | | |
|---|---|---|---|---|
| 4 | miR-143-3p | -0.470 | 0.241 | 0.782 |
| 5 | miR-22-5p | -1.303 | 0.670 | 0.782 |
| 6 | miR-141-3p | 1.723 | 0.892 | 0.782 |
| 7 | miR-133a-3p | 1.448 | 0.785 | 0.782 |
| 8 | miR-148b-3p | -0.361 | 0.197 | 0.782 |
| 9 | miR-145-5p | -1.221 | 0.680 | 0.782 |
| 10 | miR-206-3p | 1.575 | 0.901 | 0.782 |
| 11 | miR-199a-5p | 1.335 | 0.771 | 0.782 |
| 12 | miR-30c-5p | 0.438 | 0.257 | 0.782 |
| 13 | miR-451-5p | -0.638 | 0.383 | 0.782 |
| 14 | miR-16-5p | 0.649 | 0.390 | 0.782 |
| 15 | miR-181a-5p | 0.544 | 0.328 | 0.782 |
| 16 | miR-339-5p | 0.958 | 0.593 | 0.802 |
| 17 | miR-27b-3p | 0.763 | 0.515 | 0.874 |
| 18 | miR-218a-5p | -1.392 | 0.947 | 0.874 |
| 19 | miR-532-5p | 0.863 | 0.629 | 0.874 |
| 20 | miR-378a-3p | 0.664 | 0.486 | 0.874 |

miR=microRNA. Statistically significant findings are marked in bold.

**Table S3.** Twenty most significantly different miRs carried via HDL particles.

| HDL | Number | miR | log2 Fold change | SE (log Fold change) | Adjusted p-value |
|---|---|---|---|---|---|
| OVX/control | **1** | **miR-182** | -4,022 | 0,708 | **0,000** |
| vs. | **2** | **miR-125b-5p** | -2,573 | 0,446 | **0,000** |
| Sham/control | **3** | **miR-100-5p** | -3,468 | 0,634 | **0,000** |
| | **4** | **miR-221-3p** | -1,593 | 0,432 | **0,005** |
| | **5** | **miR-27b-3p** | -1,869 | 0,501 | **0,005** |
| | **6** | **miR-150-5p** | 1,336 | 0,381 | **0,008** |
| | **7** | **miR-21-5p** | -0,901 | 0,269 | **0,009** |
| | **8** | **miR-23a-3p** | -1,306 | 0,382 | **0,009** |
| | **9** | **miR-30a-5p** | -1,674 | 0,495 | **0,009** |
| | **10** | **miR-29a-3p** | -1,706 | 0,509 | **0,009** |
| | **11** | **miR-99b-5p** | -1,046 | 0,318 | **0,010** |
| | **12** | **miR-451-5p** | 1,439 | 0,470 | **0,019** |
| | **13** | **miR-103-3p** | -1,116 | 0,388 | **0,031** |
| | **14** | **miR-199a-3p** | -2,175 | 0,754 | **0,031** |
| | **15** | **miR-146b-5p** | 0,801 | 0,282 | **0,033** |
| | **16** | **miR-200b-3p** | -1,831 | 0,675 | **0,045** |
| | 17 | miR-26a-5p | 0,562 | 0,224 | 0,076 |
| | 18 | miR-152-3p | -1,255 | 0,514 | 0,086 |
| | 19 | miR-122-5p | 1,506 | 0,622 | 0,087 |
| | 20 | miR-22-3p | 0,961 | 0,423 | 0,124 |
| Sham/max | **1** | **miR-30a-5p** | -1,912 | 0,524 | **0,028** |
| vs. | **2** | **miR-100-5p** | -2,140 | 0,638 | **0,042** |
| Sham/control | 3 | miR-103-3p | -0,976 | 0,348 | 0,092 |
| | 4 | miR-27b-3p | -1,358 | 0,481 | 0,092 |
| | 5 | miR-150-5p | 1,316 | 0,471 | 0,092 |
| | 6 | miR-29a-3p | -1,484 | 0,508 | 0,092 |
| | 7 | miR-182 | -1,418 | 0,554 | 0,126 |
| | 8 | miR-378a-3p | -1,556 | 0,606 | 0,126 |
| | 9 | miR-99b-5p | -0,885 | 0,346 | 0,126 |
| | 10 | miR-184 | 1,688 | 0,675 | 0,133 |
| | 11 | miR-221-3p | -1,170 | 0,532 | 0,268 |
| | 12 | miR-30a-3p | -0,806 | 0,372 | 0,268 |
| | 13 | miR-125b-5p | -1,199 | 0,574 | 0,302 |
| | 14 | let-7b-5p | 0,487 | 0,248 | 0,381 |
| | 15 | let-7c-5p | 0,429 | 0,249 | 0,393 |
| | 16 | miR-125a-5p | -0,562 | 0,322 | 0,393 |
| | 17 | miR-146a-5p | 0,751 | 0,397 | 0,393 |
| | 18 | miR-152-3p | -0,978 | 0,579 | 0,393 |
| | 19 | miR-1b | -1,300 | 0,718 | 0,393 |
| | 20 | miR-200b-3p | -1,111 | 0,648 | 0,393 |
| OVX/max | **1** | **miR-200b-3p** | -3,439 | 0,696 | **0,000** |
| vs. | 2 | miR-206-3p | 2,394 | 0,805 | 0,097 |

| | | | | | |
|---|---|---|---|---|---|
| OVX/control | 3 | miR-155-5p | -2,302 | 0,780 | 0,097 |
| | 4 | miR-122-5p | -1,443 | 0,496 | 0,097 |
| | 5 | miR-143-3p | -0,943 | 0,340 | 0,102 |
| | 6 | miR-150-5p | -1,430 | 0,518 | 0,102 |
| | 7 | miR-342-5p | 0,838 | 0,328 | 0,148 |
| | 8 | miR-192-5p | -1,769 | 0,715 | 0,148 |
| | 9 | miR-223-5p | 0,524 | 0,213 | 0,148 |
| | 10 | miR-27a-3p | -0,493 | 0,200 | 0,148 |
| | 11 | miR-22-3p | -0,927 | 0,395 | 0,184 |
| | 12 | miR-7a-5p | 0,591 | 0,256 | 0,184 |
| | 13 | let-7g-5p | -0,497 | 0,240 | 0,313 |
| | 14 | miR-674-3p | -0,653 | 0,330 | 0,364 |
| | 15 | miR-184 | 1,659 | 0,858 | 0,370 |
| | 16 | miR-125a-5p | 0,626 | 0,328 | 0,370 |
| | 17 | miR-99a-5p | -0,431 | 0,232 | 0,370 |
| | 18 | miR-100-5p | -0,963 | 0,522 | 0,370 |
| | 19 | let-7d-3p | 0,377 | 0,207 | 0,370 |
| | 20 | miR-26b-5p | -0,835 | 0,459 | 0,370 |

miR=microRNA. Statistically significant findings are marked in bold.

**Table S4.** Twenty most significantly different rDRs carried via EVs.

| | Number | rDR | Annotation | log2 Fold change | SE (log Fold change) | Adjusted p-value |
|---|---|---|---|---|---|---|
| OVX/control | 1 | URS00008CF116_10116 | SSU rRNA | -0,788 | 0,231 | 0,068 |
| vs. | 2 | URS000069EDAA_10116 | 5S rRNA | 0,755 | 0,281 | 0,255 |
| sham/control | 3 | URS00008C5677_10116 | SSU rRNA | -0,883 | 0,325 | 0,319 |
| | 4 | URS00008C59CC_10116 | SSU rRNA | -0,419 | 0,152 | 0,332 |
| | 5 | URS00008C6042_10116 | SSU rRNA | -0,429 | 0,158 | 0,332 |
| | 6 | URS00008C644C_10116 | rRNA | -0,945 | 0,318 | 0,453 |
| | 7 | URS00008C96FC_10116 | SSU rRNA | -1,202 | 0,441 | 0,453 |
| | 8 | URS00008D1F6F_10116 | SSU rRNA | -1,101 | 0,365 | 0,617 |
| | 9 | URS000062A825_10116 | 5S rRNA | 0,674 | 0,315 | 0,643 |
| | 10 | URS00006600A3_10116 | 5S rRNA | 0,662 | 0,312 | 0,643 |
| | 11 | URS00006684C0_10116 | 5S rRNA | 0,692 | 0,306 | 0,643 |
| | 12 | URS000066C2F6_10116 | 5S rRNA | 0,647 | 0,305 | 0,664 |
| | 13 | URS00006851EC_10116 | 5S rRNA | 0,701 | 0,278 | 0,664 |
| | 14 | URS000068E3E4_10116 | 5.8S rRNA | 0,780 | 0,338 | 0,664 |
| | 15 | URS000068F5BD_10116 | 5S rRNA | 0,706 | 0,288 | 0,664 |
| | 16 | URS000069C4F1_10116 | 5S rRNA | 0,763 | 0,301 | 0,664 |
| | 17 | URS00006F3455_10116 | 5S rRNA | 0,678 | 0,294 | 0,664 |
| | 18 | URS0000706122_10116 | 5S rRNA | 0,609 | 0,282 | 0,664 |
| | 19 | URS0000706CA2_10116 | 5S rRNA | 0,681 | 0,314 | 0,664 |
| | 20 | URS00007131E0_10116 | 5S rRNA | 0,676 | 0,318 | 0,664 |
| Sham/max | 1 | URS000065B5CF_10116 | 5S rRNA (multiple genes) | -0,995 | 0,308 | 0,900 |
| vs. | 2 | URS00003F43F2_10116 | 5S RNA (Rn5s) | -0,934 | 0,316 | 0,900 |
| Sham/control | 3 | URS0000CD0331_10116 | 5S RNA (Rn5s) | -0,957 | 0,333 | 0,900 |
| | 4 | URS00006E79BE_10116 | 5.8S rRNA | -0,943 | 0,383 | 0,900 |
| | 5 | URS00006EBFDA_10116 | 5.8S rRNA | -0,915 | 0,352 | 0,900 |
| | 6 | URS00008C6266_10116 | SSU rRNA | 0,924 | 0,348 | 0,900 |
| | 7 | URS00008D160B_10116 | SSU rRNA | 0,816 | 0,315 | 0,900 |
| | 8 | URS00022B3C24_10116 | 5S rRNA | -0,893 | 0,357 | 0,900 |
| | 9 | URS00008D1F6F_10116 | SSU rRNA | 0,935 | 0,389 | 0,900 |
| | 10 | URS00022B91B4_10116 | 5S rRNA | -0,907 | 0,381 | 0,900 |
| | 11 | URS00000F9D45_10116 | RNA (121-MER)(PDB 7QGG, chain E) | -0,915 | 0,484 | 0,900 |
| | 12 | URS0000184DE3_10116 | 5S rRNA | -0,979 | 0,453 | 0,900 |
| | 13 | URS00001F5E2D_10116 | Eukaryotic LSU rRNA | 0,393 | 0,214 | 0,900 |
| | 14 | URS0000246A92_10116 | SSU rRNA | 0,387 | 0,214 | 0,900 |
| | 15 | URS000032987A_10116 | Partial rRNA | -0,508 | 0,298 | 0,900 |
| | 16 | URS000051C7BA_10116 | Eukaryotic LSU rRNA | 0,276 | 0,162 | 0,900 |
| | 17 | URS00005296BE_10116 | Miscellaneous RNA | -0,927 | 0,445 | 0,900 |
| | 18 | URS0000629A3C_10116 | 5.8S rRNA | -0,722 | 0,434 | 0,900 |
| | 19 | URS000062C46B_10116 | 5.8S rRNA | -0,539 | 0,283 | 0,900 |
| | 20 | URS0000637711_10116 | 5S rRNA | -0,719 | 0,414 | 0,900 |
| OVX/max | **1** | **URS00008C96FC_10116** | SSU rRNA | 0,930 | 0,276 | **0,011** |

| | | | | | | |
|---|---|---|---|---|---|---|
| vs. | **2** | **URS000068F5BD_10116** | 5S rRNA | -0,758 | 0,274 | **0,031** |
| OVX/control | **3** | **URS00006F129F_10116** | 5S rRNA | -0,677 | 0,267 | **0,031** |
| | 4 | URS0000706122_10116 | 5S rRNA | -0,798 | 0,300 | 0,051 |
| | 5 | URS00008C5677_10116 | SSU rRNA | 0,701 | 0,271 | 0,143 |
| | 6 | URS00008D1F6F_10116 | SSU rRNA | 0,771 | 0,303 | 0,143 |
| | 7 | URS00022B53F0_10116 | 5S rRNA | -0,759 | 0,288 | 0,157 |
| | 8 | URS00022B8935_10116 | 5S rRNA | -0,768 | 0,289 | 0,165 |
| | 9 | URS000012BFBF_10116 | SSU rRNA | 0,452 | 0,186 | 0,174 |
| | 10 | URS0000548AD6_10116 | 12S rRNA | -0,964 | 0,392 | 0,279 |
| | 11 | URS00006E79BE_10116 | 5.8S rRNA | -0,795 | 0,333 | 0,290 |
| | 12 | URS0000647009_10116 | 5S rRNA | -0,684 | 0,290 | 0,328 |
| | 13 | URS00005296BE_10116 | Miscellaneous RNA | 1,597 | 0,689 | 0,328 |
| | 14 | URS00006E6F77_10116 | 5S rRNA (multiple genes) | -0,667 | 0,299 | 0,373 |
| | 15 | URS00008CB5BD_10116 | SSU rRNA | 0,428 | 0,196 | 0,419 |
| | 16 | URS00022B6B03_10116 | 5S rRNA | -0,590 | 0,272 | 0,450 |
| | 17 | URS0000693ECA_10116 | 5S rRNA | -0,573 | 0,273 | 0,564 |
| | 18 | URS00001F5E2D_10116 | Eukaryotic LSU rRNA | 0,439 | 0,229 | 0,585 |
| | 19 | URS000064596E_10116 | 5S rRNA | -0,572 | 0,296 | 0,782 |
| | 20 | URS000065DC7C_10116 | 5S rRNA | -0,626 | 0,309 | 0,785 |

rDR=ribosomal RNA (rRNA) -derived sRNAs. Statistically significant findings are marked in bold.

| EV | Number | rDR | Annotation | log2 Fold change | SE (log Fold change) | Adjusted p-value |
|---|---|---|---|---|---|---|
| OVX/control | 1 | URS00008CF116_10116 | SSU rRNA | -0,788 | 0,231 | 0,130 |
| vs. | 2 | URS000069EDAA_10116 | 5S rRNA | 0,755 | 0,281 | 0,185 |
| Sham/control | 3 | URS00008C5677_10116 | SSU rRNA | -0,883 | 0,325 | 0,185 |
| | 4 | URS00008C59CC_10116 | SSU rRNA | -0,419 | 0,152 | 0,185 |
| | 5 | URS00008C6042_10116 | SSU rRNA | -0,429 | 0,158 | 0,185 |
| | 6 | URS00008C644C_10116 | rRNA | -0,945 | 0,318 | 0,185 |
| | 7 | URS00008C96FC_10116 | SSU rRNA | -1,202 | 0,441 | 0,185 |
| | 8 | URS00008D1F6F_10116 | SSU rRNA | -1,101 | 0,365 | 0,185 |
| | 9 | URS000062A825_10116 | 5S rRNA | 0,674 | 0,315 | 0,219 |
| | 10 | URS00006600A3_10116 | 5S rRNA | 0,662 | 0,312 | 0,219 |
| | 11 | URS00006684C0_10116 | 5S rRNA | 0,692 | 0,306 | 0,219 |
| | 12 | URS000066C2F6_10116 | 5S rRNA | 0,647 | 0,305 | 0,219 |
| | 13 | URS00006851EC_10116 | 5S rRNA | 0,701 | 0,278 | 0,219 |
| | 14 | URS000068E3E4_10116 | 5.8S rRNA | 0,780 | 0,338 | 0,219 |
| | 15 | URS000068F5BD_10116 | 5S rRNA | 0,706 | 0,288 | 0,219 |
| | 16 | URS000069C4F1_10116 | 5S rRNA | 0,763 | 0,301 | 0,219 |
| | 17 | URS00006F3455_10116 | 5S rRNA | 0,678 | 0,294 | 0,219 |
| | 18 | URS0000706122_10116 | 5S rRNA | 0,609 | 0,282 | 0,219 |
| | 19 | URS0000706CA2_10116 | 5S rRNA | 0,681 | 0,314 | 0,219 |
| | 20 | URS00007131E0_10116 | 5S rRNA | 0,676 | 0,318 | 0,219 |
| Sham/max | 1 | URS000065B5CF_10116 | 5S rRNA (multiple genes) | -0,995 | 0,308 | 0,283 |
| vs. | 2 | URS00003F43F2_10116 | 5S RNA (Rn5s) | -0,934 | 0,316 | 0,307 |
| Sham/control | 3 | URS0000CD0331_10116 | 5S RNA (Rn5s) | -0,957 | 0,333 | 0,307 |
| | 4 | URS00006E79BE_10116 | 5.8S rRNA | -0,943 | 0,383 | 0,323 |
| | 5 | URS00006EBFDA_10116 | 5.8S rRNA | -0,915 | 0,352 | 0,323 |
| | 6 | URS00008C6266_10116 | SSU rRNA | 0,924 | 0,348 | 0,323 |

| | | | | | | |
|---|---|---|---|---|---|---|
| | 7 | URS00008D160B_10116 | SSU rRNA | 0,816 | 0,315 | 0,323 |
| | 8 | URS00022B3C24_10116 | 5S rRNA | -0,893 | 0,357 | 0,323 |
| | 9 | URS00008D1F6F_10116 | SSU rRNA | 0,935 | 0,389 | 0,356 |
| | 10 | URS00022B91B4_10116 | 5S rRNA | -0,907 | 0,381 | 0,356 |
| | 11 | URS00000F9D45_10116 | RNA (121-MER)(PDB 7QGG, chain E) | -0,915 | 0,484 | 0,365 |
| | 12 | URS0000184DE3_10116 | 5S rRNA | -0,979 | 0,453 | 0,365 |
| | 13 | URS00001F5E2D_10116 | Eukaryotic LSU rRNA | 0,393 | 0,214 | 0,365 |
| | 14 | URS0000246A92_10116 | SSU rRNA | 0,387 | 0,214 | 0,365 |
| | 15 | URS000032987A_10116 | Partial rRNA | -0,508 | 0,298 | 0,365 |
| | 16 | URS000051C7BA_10116 | Eukaryotic LSU rRNA | 0,276 | 0,162 | 0,365 |
| | 17 | URS00005296BE_10116 | Miscellaneous RNA | -0,927 | 0,445 | 0,365 |
| | 18 | URS0000629A3C_10116 | 5.8S rRNA | -0,722 | 0,434 | 0,365 |
| | 19 | URS000062C46B_10116 | 5.8S rRNA | -0,539 | 0,283 | 0,365 |
| | 20 | URS0000637711_10116 | 5S rRNA | -0,719 | 0,414 | 0,365 |
| OVX/max vs. OVX/control | 1 | URS00008C96FC_10116 | SSU rRNA | 0,930 | 0,276 | 0,156 |
| | 2 | URS000068F5BD_10116 | 5S rRNA | -0,758 | 0,274 | 0,285 |
| | 3 | URS00006F129F_10116 | 5S rRNA | -0,677 | 0,267 | 0,285 |
| | 4 | URS0000706122_10116 | 5S rRNA | -0,798 | 0,300 | 0,285 |
| | 5 | URS00008C5677_10116 | SSU rRNA | 0,701 | 0,271 | 0,285 |
| | 6 | URS00008D1F6F_10116 | SSU rRNA | 0,771 | 0,303 | 0,285 |
| | 7 | URS00022B53F0_10116 | 5S rRNA | -0,759 | 0,288 | 0,285 |
| | 8 | URS00022B8935_10116 | 5S rRNA | -0,768 | 0,289 | 0,285 |
| | 9 | URS000012BFBF_10116 | SSU rRNA | 0,452 | 0,186 | 0,314 |
| | 10 | URS0000548AD6_10116 | 12S rRNA | -0,964 | 0,392 | 0,314 |
| | 11 | URS00006E79BE_10116 | 5.8S rRNA | -0,795 | 0,333 | 0,314 |
| | 12 | URS0000647009_10116 | 5S rRNA | -0,684 | 0,290 | 0,316 |
| | 13 | URS00005296BE_10116 | Miscellaneous RNA | 1,597 | 0,689 | 0,325 |
| | 14 | URS00006E6F77_10116 | 5S rRNA (multiple genes) | -0,667 | 0,299 | 0,383 |
| | 15 | URS00008CB5BD_10116 | SSU rRNA | 0,428 | 0,196 | 0,388 |
| | 16 | URS00022B6B03_10116 | 5S rRNA | -0,590 | 0,272 | 0,388 |
| | 17 | URS0000693ECA_10116 | 5S rRNA | -0,573 | 0,273 | 0,432 |
| | 18 | URS00001F5E2D_10116 | Eukaryotic LSU rRNA | 0,439 | 0,229 | 0,448 |
| | 19 | URS000064596E_10116 | 5S rRNA | -0,572 | 0,296 | 0,448 |
| | 20 | URS000065DC7C_10116 | 5S rRNA | -0,626 | 0,309 | 0,448 |

rDR=ribosomal RNA (rRNA) -derived sRNAs. Statistically significant findings are marked in bold.

**Table S5.** Twenty most significantly different rDRs carried via HDL particles.

| | Number | rDR | Annotation | log2 Fold change | SE (log Fold change) | Adjusted p-value |
|---|---|---|---|---|---|---|
| OVX/control | 1 | URS0001B4B318_10116 | SSU rRNA | 1,929 | 0,548 | 0,0675745 |
| vs. | 2 | URS0001B0949E_10116 | SSU rRNA | 1,701 | 0,579 | 0,254542 |
| Sham/control | 3 | URS000065DC7C_10116 | 5S rRNA | 1,715 | 0,626 | 0,3189266 |
| | 4 | URS00008CB13B_10116 | SSU rRNA | 1,677 | 0,645 | 0,3324486 |
| | 5 | URS00022B0F36_10116 | 5S rRNA | 1,292 | 0,506 | 0,3324486 |
| | 6 | URS0000623A69_10116 | 5S rRNA | 1,238 | 0,521 | 0,4530505 |
| | 7 | URS00006EFC8C_10116 | 5S rRNA | 1,639 | 0,707 | 0,4530505 |
| | 8 | URS000069C4F1_10116 | 5S rRNA | 0,986 | 0,459 | 0,6173743 |
| | 9 | URS000066C2F6_10116 | 5S rRNA | 0,891 | 0,438 | 0,6429183 |
| | 10 | URS00006CE1FB_10116 | 5.8S rRNA | 0,974 | 0,487 | 0,6429183 |
| | 11 | URS00008C5C0D_10116 | SSU rRNA | 0,674 | 0,331 | 0,6429183 |
| | 12 | URS0000005270_10116 | RNA (157-MER)(PDB 7QGG, chain D) | 0,862 | 0,492 | 0,6641598 |
| | 13 | URS0000070986_10116 | Partial rRNA | 0,856 | 0,722 | 0,6641598 |
| | 14 | URS00001C1BDA_10116 | 28S rRNA | 0,438 | 0,300 | 0,6641598 |
| | 15 | URS0000315E21_10116 | rRNA | 0,850 | 0,499 | 0,6641598 |
| | 16 | URS00003E4DE0_10116 | Eukaryotic LSU rRNA | 0,569 | 0,444 | 0,6641598 |
| | 17 | URS00003ECA53_10116 | Eukaryotic LSU rRNA | -0,434 | 0,379 | 0,6641598 |
| | 18 | URS00004213B9_10116 | Partial rRNA | 0,805 | 0,679 | 0,6641598 |
| | 19 | URS00004B2C76_10116 | 5.8S rRNA (Rn5-8s) | 0,864 | 0,499 | 0,6641598 |
| | 20 | URS00005296BE_10116 | Miscellaneous RNA | 0,958 | 0,677 | 0,6641598 |
| Sham/max | 1 | URS0000005270_10116 | RNA (157-MER)(PDB 7QGG, chain D) | 0,211 | 0,322 | 0,9002191 |
| vs. | 2 | URS00000351A9_10116 | 16S rRNA | 0,248 | 0,419 | 0,9002191 |
| Sham/control | 3 | URS000006A105_10116 | 16S rRNA | 0,430 | 0,415 | 0,9002191 |
| | 4 | URS0000070986_10116 | Partial rRNA | -0,242 | 0,423 | 0,9002191 |
| | 5 | URS00000F446C_10116 | Partial rRNA | -0,262 | 0,303 | 0,9002191 |
| | 6 | URS00000F8D11_10116 | Partial 18S rRNA | -0,236 | 0,336 | 0,9002191 |
| | 7 | URS00000F9D45_10116 | RNA (121-MER)(PDB 7QGG, chain E) | -0,268 | 0,422 | 0,9002191 |
| | 8 | URS0000114737_10116 | 16S rRNA | -0,403 | 0,462 | 0,9002191 |
| | 9 | URS0000121DBD_10116 | Partial rRNA | -0,310 | 0,292 | 0,9002191 |
| | 10 | URS0000129B9F_10116 | 5S rRNA (variant) | -0,267 | 0,389 | 0,9002191 |
| | 11 | URS00001C1BDA_10116 | 28S rRNA | 0,323 | 0,263 | 0,9002191 |
| | 12 | URS00001CAE00_10116 | rRNA | 0,347 | 0,324 | 0,9002191 |
| | 13 | URS00001F5E2D_10116 | Eukaryotic LSU rRNA | 0,269 | 0,217 | 0,9002191 |
| | 14 | URS0000246A92_10116 | SSU rRNA | -0,223 | 0,344 | 0,9002191 |
| | 15 | URS0000315E21_10116 | rRNA | 0,204 | 0,328 | 0,9002191 |
| | 16 | URS000032987A_10116 | Partial rRNA | 0,225 | 0,384 | 0,9002191 |
| | 17 | URS00003C1820_10116 | 12S rRNA | 0,704 | 0,512 | 0,9002191 |
| | 18 | URS00003E4DE0_10116 | Eukaryotic LSU rRNA | -0,154 | 0,275 | 0,9002191 |
| | 19 | URS00003ECA53_10116 | Eukaryotic LSU rRNA | -0,289 | 0,345 | 0,9002191 |
| | 20 | URS00003F43F2_10116 | 5S RNA (Rn5s) | -0,488 | 0,444 | 0,9002191 |

| | | | | | | |
|---|---|---|---|---|---|---|
| OVX/max vs. OVX/control | **1** | **URS00008CB13B_10116** | SSU rRNA | -2,290 | 0,576 | **0,0107535** |
| | **2** | **URS0001B0949E_10116** | SSU rRNA | -1,863 | 0,526 | **0,0308956** |
| | **3** | **URS0001B4B318_10116** | SSU rRNA | -1,820 | 0,530 | **0,0308956** |
| | 4 | URS00008C5C0D_10116 | SSU rRNA | -0,900 | 0,280 | 0,0509012 |
| | 5 | URS00008C5677_10116 | SSU rRNA | -0,853 | 0,303 | 0,1432148 |
| | 6 | URS00008C96FC_10116 | SSU rRNA | -0,865 | 0,312 | 0,1432148 |
| | 7 | URS00008D1F6F_10116 | SSU rRNA | -0,879 | 0,326 | 0,1573582 |
| | 8 | URS00008C644C_10116 | SSU rRNA | 0,801 | 0,305 | 0,1654317 |
| | 9 | URS00008C6266_10116 | SSU rRNA | -0,833 | 0,324 | 0,1743792 |
| | 10 | URS000012BFBF_10116 | SSU rRNA | -0,524 | 0,222 | 0,2794299 |
| | 11 | URS00008CAD43_10116 | SSU rRNA | -0,829 | 0,358 | 0,2900912 |
| | 12 | URS00005296BE_10116 | Miscellaneous RNA | -1,356 | 0,615 | 0,3279632 |
| | 13 | URS00008D160B_10116 | SSU rRNA | -0,795 | 0,358 | 0,3279632 |
| | 14 | URS000051C7BA_10116 | Eukaryotic LSU rRNA | -0,539 | 0,254 | 0,3727819 |
| | 15 | URS00008C8B6E_10116 | SSU rRNA | 0,319 | 0,156 | 0,4194062 |
| | 16 | URS00009365D8_10116 | Eukaryotic large subunit ribosomal RNA | -1,215 | 0,610 | 0,4502202 |
| | 17 | URS00008C6042_10116 | SSU rRNA | 0,304 | 0,163 | 0,5636067 |
| | 18 | URS00008C47B3_10116 | SSU rRNA | -0,614 | 0,336 | 0,5850909 |
| | 19 | URS00008C6B45_10116 | SSU rRNA | -0,221 | 0,133 | 0,7823364 |
| | 20 | URS00000F446C_10116 | Partial rRNA | 0,376 | 0,229 | 0,7854258 |

rDR=ribosomal RNA (rRNA) -derived sRNA. SSU=small subunit, LSU=large subunit. Statistically significant findings are marked in bold.

**Table S6.** Twenty most significantly different tDRs carried via EVs.

| **EV** | **Number** | **tDR** | **log2 Fold change** | **SE (log Fold change)** | **Adjusted p-value** |
|---|---|---|---|---|---|
| OVX/control | **1** | **tRNA-Met-CAT-3-1** | 4,196 | 0,699 | **0,000** |
| vs. | **2** | **tRNA-Met-CAT-1-1** | 3,361 | 0,652 | **0,000** |
| Sham/control | 3 | tRNA-Ala-TGC-1-1 | 0,799 | 0,412 | 0,654 |
| | 4 | tRNA-Ala-TGC-1-2 | 0,733 | 0,455 | 0,654 |
| | 5 | tRNA-Arg-CCT-3-1 | 1,101 | 0,565 | 0,654 |
| | 6 | tRNA-Arg-CCT-4-1 | 1,073 | 0,565 | 0,654 |
| | 7 | tRNA-Asp-GTC-1-1 | -0,997 | 0,430 | 0,654 |
| | 8 | tRNA-Glu-CTC-1-10 | -0,712 | 0,407 | 0,654 |
| | 9 | tRNA-Glu-CTC-1-11 | -0,711 | 0,412 | 0,654 |
| | 10 | tRNA-Glu-CTC-1-12 | -0,674 | 0,414 | 0,654 |
| | 11 | tRNA-Glu-CTC-1-2 | -0,712 | 0,410 | 0,654 |
| | 12 | tRNA-Glu-CTC-1-3 | -0,684 | 0,406 | 0,654 |
| | 13 | tRNA-Glu-CTC-1-4 | -0,689 | 0,411 | 0,654 |
| | 14 | tRNA-Glu-CTC-1-7 | -0,690 | 0,413 | 0,654 |
| | 15 | tRNA-Glu-CTC-1-9 | -0,729 | 0,415 | 0,654 |
| | 16 | tRNA-Glu-CTC-2-1 | 0,525 | 0,296 | 0,654 |
| | 17 | tRNA-Glu-CTC-4-1 | -0,678 | 0,409 | 0,654 |
| | 18 | tRNA-Glu-TTC-2-1 | -0,588 | 0,324 | 0,654 |
| | 19 | tRNA-Glu-TTC-3-1 | -0,484 | 0,303 | 0,654 |
| | 20 | tRNA-Lys-CTT-1-4 | -0,630 | 0,313 | 0,654 |
| Sham/max | **1** | **tRNA-Met-CAT-3-1** | 3,802 | 0,706 | **0,000** |
| vs. | 2 | tRNA-Leu-AAG-2-1 | 2,458 | 0,739 | 0,058 |
| Sham/control | 3 | tRNA-Met-CAT-1-1 | 2,418 | 0,734 | 0,058 |
| | 4 | tRNA-Leu-AAG-2-2 | 2,231 | 0,734 | 0,084 |
| | 5 | tRNA-Leu-AAG-2-3 | 2,275 | 0,742 | 0,084 |
| | 6 | tRNA-Glu-CTC-1-1 | -0,887 | 0,419 | 0,332 |
| | 7 | tRNA-Glu-CTC-1-10 | -0,924 | 0,418 | 0,332 |
| | 8 | tRNA-Glu-CTC-1-11 | -0,984 | 0,418 | 0,332 |
| | 9 | tRNA-Glu-CTC-1-2 | -0,875 | 0,416 | 0,332 |
| | 10 | tRNA-Glu-CTC-1-3 | -0,914 | 0,418 | 0,332 |
| | 11 | tRNA-Glu-CTC-1-6 | -0,886 | 0,419 | 0,332 |
| | 12 | tRNA-Glu-CTC-1-7 | -0,905 | 0,422 | 0,332 |
| | 13 | tRNA-Glu-CTC-1-8 | -0,923 | 0,426 | 0,332 |
| | 14 | tRNA-Glu-CTC-1-9 | -0,903 | 0,420 | 0,332 |
| | 15 | tRNA-Glu-CTC-4-1 | -0,901 | 0,417 | 0,332 |
| | 16 | tRNA-Leu-CAA-1-1 | 1,311 | 0,571 | 0,332 |
| | 17 | tRNA-Leu-CAA-2-1 | 1,355 | 0,566 | 0,332 |
| | 18 | tRNA-Ser-AGA-3-1 | 1,323 | 0,531 | 0,332 |
| | 19 | tRNA-Ser-GCT-3-4 | 1,189 | 0,534 | 0,332 |
| | 20 | tRNA-Glu-CTC-1-12 | -0,874 | 0,420 | 0,332 |
| OVX/max | 1 | tRNA-Ala-CGC-3-1 | 0,953 | 0,414 | 0,632 |

| | | | | | |
|---|---|---|---|---|---|
| vs. | 2 | tRNA-Glu-TTC-2-2 | -0,575 | 0,244 | 0,632 |
| OVX/control | 3 | tRNA-Glu-TTC-3-1 | -0,479 | 0,201 | 0,632 |
| | 4 | tRNA-Gly-CCC-3-1 | -1,082 | 0,459 | 0,632 |
| | 5 | tRNA-Lys-TTT-4-1 | 1,633 | 0,636 | 0,632 |
| | 6 | tRNA-Pro-CGG-1-1 | 0,526 | 0,238 | 0,632 |
| | 7 | tRNA-Pro-TGG-2-1 | 1,161 | 0,530 | 0,632 |
| | 8 | tRNA-SeC-TCA-1-1 | -1,431 | 0,526 | 0,632 |
| | 9 | tRNA-Ala-AGC-1-1 | 0,800 | 0,433 | 0,669 |
| | 10 | tRNA-Ala-AGC-2-1 | 0,783 | 0,553 | 0,669 |
| | 11 | tRNA-Ala-AGC-2-2 | 0,832 | 0,555 | 0,669 |
| | 12 | tRNA-Ala-AGC-3-1 | 0,676 | 0,546 | 0,669 |
| | 13 | tRNA-Ala-AGC-3-3 | 0,879 | 0,550 | 0,669 |
| | 14 | tRNA-Ala-CGC-1-1 | 0,857 | 0,469 | 0,669 |
| | 15 | tRNA-Ala-CGC-2-1 | 0,670 | 0,403 | 0,669 |
| | 16 | tRNA-Ala-TGC-1-1 | 0,464 | 0,426 | 0,669 |
| | 17 | tRNA-Ala-TGC-2-1 | 0,484 | 0,402 | 0,669 |
| | 18 | tRNA-Ala-TGC-3-1 | 0,811 | 0,556 | 0,669 |
| | 19 | tRNA-Ala-TGC-4-1 | 0,727 | 0,588 | 0,669 |
| | 20 | tRNA-Arg-CCT-3-1 | -0,433 | 0,414 | 0,669 |

tDR=transfer RNA (rRNA) -derived sRNAs. Statistically significant findings are marked in bold.

**Table S7.** Twenty most significantly different tDRs carried via HDL particles.

| HDL | Number | tDR | log2 Fold change | SE (log Fold change) | Adjusted p-value |
|---|---|---|---|---|---|
| OVX/control | 1 | tRNA-Glu-TTC-2-2 | 1,012 | 0,411 | 0,306 |
| vs. | 2 | tRNA-Gly-GCC-2-1 | -0,426 | 0,167 | 0,306 |
| Sham/control | 3 | tRNA-Gly-GCC-2-6 | -0,398 | 0,153 | 0,306 |
| | 4 | tRNA-Gly-GCC-4-1 | -0,383 | 0,131 | 0,306 |
| | 5 | tRNA-His-GTG-1-2 | -1,158 | 0,478 | 0,306 |
| | 6 | tRNA-His-GTG-1-3 | -1,195 | 0,466 | 0,306 |
| | 7 | tRNA-His-GTG-1-5 | -1,211 | 0,484 | 0,306 |
| | 8 | tRNA-His-GTG-1-8 | -1,211 | 0,472 | 0,306 |
| | 9 | tRNA-Ala-AGC-1-1 | 1,518 | 0,706 | 0,425 |
| | 10 | tRNA-Glu-TTC-2-1 | 0,868 | 0,403 | 0,425 |
| | 11 | tRNA-Glu-TTC-3-1 | 0,887 | 0,414 | 0,425 |
| | 12 | tRNA-Gly-GCC-2-2 | -0,367 | 0,165 | 0,425 |
| | 13 | tRNA-Gly-GCC-2-3 | -0,354 | 0,168 | 0,425 |
| | 14 | tRNA-Gly-GCC-2-4 | -0,383 | 0,190 | 0,440 |
| | 15 | tRNA-Gly-GCC-2-5 | -0,331 | 0,165 | 0,440 |
| | 16 | tRNA-His-GTG-1-10 | -1,049 | 0,519 | 0,440 |
| | 17 | tRNA-His-GTG-1-7 | -0,964 | 0,485 | 0,440 |
| | 18 | tRNA-His-GTG-1-11 | -0,934 | 0,481 | 0,443 |
| | 19 | tRNA-Pro-CGG-1-1 | -0,786 | 0,406 | 0,443 |
| | 20 | tRNA-Gln-CTG-1-1 | 0,997 | 0,538 | 0,509 |
| Sham/max | 1 | tRNA-Arg-CCG-3-1 | -3,095 | 0,991 | 0,265 |
| vs. | 2 | tRNA-Pro-CGG-1-1 | -1,281 | 0,437 | 0,265 |
| Sham/control | 3 | tRNA-Ala-AGC-1-1 | 1,261 | 0,701 | 0,968 |
| | 4 | tRNA-Ala-AGC-2-2 | 0,465 | 0,576 | 0,968 |
| | 5 | tRNA-Ala-AGC-3-1 | 0,555 | 0,605 | 0,968 |
| | 6 | tRNA-Ala-AGC-3-3 | 0,658 | 0,640 | 0,968 |
| | 7 | tRNA-Ala-CGC-1-1 | 0,971 | 0,743 | 0,968 |
| | 8 | tRNA-Ala-CGC-2-1 | 0,599 | 0,678 | 0,968 |
| | 9 | tRNA-Ala-CGC-3-1 | 0,621 | 0,713 | 0,968 |
| | 10 | tRNA-Ala-TGC-1-1 | 0,614 | 0,690 | 0,968 |
| | 11 | tRNA-Ala-TGC-1-2 | 0,696 | 0,671 | 0,968 |
| | 12 | tRNA-Ala-TGC-2-1 | 0,882 | 0,798 | 0,968 |
| | 13 | tRNA-Ala-TGC-3-1 | 0,482 | 0,597 | 0,968 |
| | 14 | tRNA-Ala-TGC-4-1 | 0,726 | 0,751 | 0,968 |
| | 15 | tRNA-Asp-GTC-2-11 | 0,437 | 0,534 | 0,968 |
| | 16 | tRNA-Asp-GTC-2-14 | 0,586 | 0,650 | 0,968 |
| | 17 | tRNA-Asp-GTC-2-5 | 0,552 | 0,595 | 0,968 |
| | 18 | tRNA-Cys-GCA-1-1 | 0,414 | 0,488 | 0,968 |
| | 19 | tRNA-Gln-CTG-1-1 | 0,481 | 0,448 | 0,968 |
| | 20 | tRNA-Gln-CTG-1-2 | 0,437 | 0,462 | 0,968 |
| OVX/max | 1 | tRNA-Leu-AAG-2-1 | -1,988 | 0,639 | 0,297 |
| vs. | 2 | tRNA-Leu-AAG-2-3 | -1,714 | 0,594 | 0,309 |
| OVX/control | 3 | tRNA-Gln-CTG-7-1 | -1,511 | 0,598 | 0,608 |
| | 4 | tRNA-Ala-AGC-1-1 | -0,524 | 0,571 | 0,987 |
| | 5 | tRNA-Ala-CGC-1-1 | -0,492 | 0,565 | 0,987 |
| | 6 | tRNA-Ala-CGC-2-1 | -0,957 | 0,560 | 0,987 |
| | 7 | tRNA-Ala-CGC-3-1 | -0,298 | 0,522 | 0,987 |
| | 8 | tRNA-Ala-TGC-1-1 | -0,891 | 0,610 | 0,987 |
| | 9 | tRNA-Ala-TGC-1-2 | -0,942 | 0,608 | 0,987 |

| | | | | |
|---|---|---|---|---|
| 10 | tRNA-Ala-TGC-2-1 | -0,612 | 0,583 | 0,987 |
| 11 | tRNA-Ala-TGC-3-1 | 0,292 | 0,501 | 0,987 |
| 12 | tRNA-Ala-TGC-4-1 | -0,711 | 0,619 | 0,987 |
| 13 | tRNA-Arg-CCG-3-1 | -0,875 | 0,720 | 0,987 |
| 14 | tRNA-Asp-GTC-2-2 | 0,417 | 0,439 | 0,987 |
| 15 | tRNA-Cys-GCA-11-1 | -0,924 | 0,567 | 0,987 |
| 16 | tRNA-Cys-GCA-1-3 | 0,309 | 0,420 | 0,987 |
| 17 | tRNA-Cys-GCA-1-4 | 0,459 | 0,438 | 0,987 |
| 18 | tRNA-Cys-GCA-2-1 | 0,406 | 0,454 | 0,987 |
| 19 | tRNA-Cys-GCA-2-2 | 0,554 | 0,472 | 0,987 |
| 20 | tRNA-Gln-CTG-1-1 | -0,426 | 0,423 | 0,987 |

tDR=transfer RNA (rRNA) -derived sRNAs. Statistically significant findings are marked in bold.

**Table S8.** Significant Kyoto Encyclopedia of Genes and Genomes (KEGG) pathways of significantly different miRs carried via EVs.

| | Number | KEGG pathway | p-value | number of regulated genes | number of regulating miRs |
|---|---|---|---|---|---|
| OVX/control vs. | 1 | Proteoglycans in cancer | 0.038 | 2 | 1 |
| Sham/control | 2 | Type II diabetes mellitus | 0.057 | 4 | 3 |
| Sham/max | 1 | Adipocytokine signaling pathway | 0.025 | 1 | 1 |
| vs. | 2 | ErbB signaling pathway | 0.030 | 1 | 1 |
| Sham/control | 3 | Wnt signaling pathway | 0.032 | 1 | 1 |
| | 4 | MicroRNAs in cancer | 0.035 | 2 | 1 |
| | 5 | HTLV-I infection | 0.039 | 1 | 1 |
| | 6 | Caffeine metabolism | 0.039 | 1 | 1 |
| | 7 | Pathways in cancer | 0.040 | 3 | 1 |
| | 8 | Proteoglycans in cancer | 0.046 | 2 | 1 |
| | 9 | MAPK signaling pathway | 0.051 | 1 | 1 |
| | 10 | HIF-1 signaling pathway | 0.052 | 1 | 1 |

| EV | Number | KEGG pathway (pathways union) | p-value | number of regulated genes | number of regulating miRs |
|---|---|---|---|---|---|
| OVX/control vs. | 1 | Proteoglycans in cancer | 0,038 | 2 | 1 |
| Sham/control | 2 | Type II diabetes mellitus | 0,057 | 4 | 3 |
| Sham/max | 1 | Adipocytokine signaling pathway | 0,025 | 1 | 1 |
| vs. | 2 | ErbB signaling pathway | 0,030 | 1 | 1 |
| Sham/control | 3 | Wnt signaling pathway | 0,032 | 1 | 1 |
| | 4 | MicroRNAs in cancer | 0,035 | 2 | 1 |
| | 5 | HTLV-I infection | 0,039 | 1 | 1 |
| | 6 | Caffeine metabolism | 0,040 | 1 | 1 |
| | 7 | Pathways in cancer | 0,040 | 3 | 1 |
| | 8 | Proteoglycans in cancer | 0,046 | 2 | 1 |
| | 9 | MAPK signaling pathway | 0,051 | 1 | 1 |
| | 10 | HIF-1 signaling pathway | 0,052 | 1 | 1 |

**Table S9.** Significant Kyoto Encyclopedia of Genes and Genomes (KEGG) pathways of significantly different miRs carried via HDL particles.

| | Number | KEGG pathway (pathways union) | p-value | number of regulated genes | number of regulating miRs |
|---|---|---|---|---|---|
| OVX/control | 1 | MicroRNAs in cancer | 0.003 | 2 | 2 |
| vs. | 2 | Proteoglycans in cancer | 0.014 | 2 | 2 |
| Sham/control | 3 | PI3K-Akt signaling pathway | 0.016 | 2 | 2 |
| | 4 | Pathways in cancer | 0.019 | 3 | 2 |
| | 5 | Endocytosis | 0.030 | 5 | 3 |
| | 6 | MAPK signaling pathway | 0.031 | 5 | 3 |
| | 7 | HIF-1 signaling pathway | 0.054 | 2 | 2 |
| Sham/max | 1 | Type II diabetes mellitus | 0.002 | 1 | 1 |
| vs. | 2 | Central carbon metabolism in cancer | 0.003 | 2 | 1 |
| Sham/control | 3 | Glycosaminoglycan biosynthesis - heparan sulfate / heparin | 0.003 | 1 | 1 |
| | 4 | Proteoglycans in cancer | 0.006 | 2 | 1 |
| | 5 | ErbB signaling pathway | 0.008 | 1 | 1 |
| | 6 | mTOR signaling pathway | 0.009 | 1 | 1 |
| | 7 | Adipocytokine signaling pathway | 0.010 | 1 | 1 |
| | 8 | HIF-1 signaling pathway | 0.011 | 1 | 1 |
| | 9 | Signaling pathways regulating pluripotency of stem cells | 0.013 | 2 | 1 |
| | 10 | MicroRNAs in cancer | 0.013 | 2 | 1 |
| | 11 | Glioma | 0.014 | 1 | 1 |
| | 12 | Choline metabolism in cancer | 0.016 | 1 | 1 |
| | 13 | Basal cell carcinoma | 0.017 | 1 | 1 |
| | 14 | Pathways in cancer | 0.017 | 3 | 1 |
| | 15 | Acute myeloid leukemia | 0.017 | 1 | 1 |
| | 16 | Insulin signaling pathway | 0.019 | 1 | 1 |
| | 17 | Rap1 signaling pathway | 0.020 | 1 | 1 |
| | 18 | Endocytosis | 0.022 | 1 | 1 |
| | 19 | Regulation of actin cytoskeleton | 0.023 | 1 | 1 |
| | 20 | Thyroid hormone signaling pathway | 0.024 | 1 | 1 |
| | 21 | Melanogenesis | 0.026 | 1 | 1 |
| | 22 | AMPK signaling pathway | 0.026 | 1 | 1 |
| | 23 | Bladder cancer | 0.026 | 1 | 1 |
| | 24 | HTLV-I infection | 0.030 | 1 | 1 |
| | 25 | PI3K-Akt signaling pathway | 0.030 | 2 | 1 |
| | 26 | Wnt signaling pathway | 0.031 | 1 | 1 |
| | 27 | Prostate cancer | 0.034 | 1 | 1 |
| | 28 | Ras signaling pathway | 0.042 | 1 | 1 |
| | 29 | Hippo signaling pathway | 0.044 | 1 | 1 |
| | 30 | MAPK signaling pathway | 0.047 | 1 | 1 |

| HDL | Number | KEGG pathway (pathways union) | p-value | number of regulated genes | number of regulating miRs |
|---|---|---|---|---|---|
| OVX/control | 1 | MicroRNAs in cancer | 0,003 | 2 | 2 |
| vs. | 2 | Proteoglycans in cancer | 0,014 | 2 | 2 |
| Sham/control | 3 | PI3K-Akt signaling pathway | 0,016 | 2 | 2 |
| | 4 | Pathways in cancer | 0,019 | 3 | 2 |
| | 5 | Endocytosis | 0,030 | 5 | 3 |
| | 6 | MAPK signaling pathway | 0,031 | 5 | 3 |
| | 7 | HIF-1 signaling pathway | 0,054 | 2 | 2 |
| Sham/max | 1 | Type II diabetes mellitus | 0,002 | 1 | 1 |
| vs. | 2 | Central carbon metabolism in cancer | 0,003 | 2 | 1 |
| Sham/control | 3 | Glycosaminoglycan biosynthesis - heparan sulfate / heparin | 0,003 | 1 | 1 |
| | 4 | Proteoglycans in cancer | 0,006 | 2 | 1 |
| | 5 | ErbB signaling pathway | 0,008 | 1 | 1 |
| | 6 | mTOR signaling pathway | 0,009 | 1 | 1 |
| | 7 | Adipocytokine signaling pathway | 0,010 | 1 | 1 |
| | 8 | HIF-1 signaling pathway | 0,011 | 1 | 1 |
| | 9 | Signaling pathways regulating pluripotency of stem cells | 0,013 | 2 | 1 |
| | 10 | MicroRNAs in cancer | 0,013 | 2 | 1 |
| | 11 | Glioma | 0,014 | 1 | 1 |
| | 12 | Choline metabolism in cancer | 0,016 | 1 | 1 |
| | 13 | Basal cell carcinoma | 0,017 | 1 | 1 |
| | 14 | Pathways in cancer | 0,017 | 3 | 1 |
| | 15 | Acute myeloid leukemia | 0,017 | 1 | 1 |
| | 16 | Insulin signaling pathway | 0,019 | 1 | 1 |
| | 17 | Rap1 signaling pathway | 0,020 | 1 | 1 |
| | 18 | Endocytosis | 0,022 | 1 | 1 |
| | 19 | Regulation of actin cytoskeleton | 0,023 | 1 | 1 |
| | 20 | Thyroid hormone signaling pathway | 0,024 | 1 | 1 |
| | 21 | Melanogenesis | 0,026 | 1 | 1 |
| | 22 | AMPK signaling pathway | 0,026 | 1 | 1 |
| | 23 | Bladder cancer | 0,026 | 1 | 1 |
| | 24 | HTLV-I infection | 0,030 | 1 | 1 |
| | 25 | PI3K-Akt signaling pathway | 0,030 | 2 | 1 |
| | 26 | Wnt signaling pathway | 0,031 | 1 | 1 |
| | 27 | Prostate cancer | 0,034 | 1 | 1 |
| | 28 | Ras signaling pathway | 0,042 | 1 | 1 |
| | 29 | Hippo signaling pathway | 0,044 | 1 | 1 |
| | 30 | MAPK signaling pathway | 0,047 | 1 | 1 |
